# Evolution of compound eye cell types shapes visual behaviors across *Heliconius* butterflies

**DOI:** 10.64898/2026.07.22.740151

**Authors:** Wei Lu, Nicholas W. VanKuren, Christopher Catalano, Michael S. Levine, Charlotte Wallsten, Marcus R. Kronforst

## Abstract

The evolution of visual systems is tightly linked to the diversification of visually guided behaviors. Using *Heliconius* butterflies, renowned for their rapid diversification of wing color patterns and corresponding visual mate preferences, we investigate the evolution of compound eye cell types across a complete speciation continuum. By integrating population genomics, chromatin conformation capture, and single-nucleus multi-omics, we uncover multi-modal mechanisms driving visual system evolution. Within a population polymorphic for mate preference, we identify *senseless-2* as a glia-expressing candidate visual preference gene. Across species, we reveal that photoreceptors, particularly the color-sensing R7s, are the fastest-evolving cell types. Furthermore, we molecularly characterize novel R2/5 and R7 photoreceptor subtypes, showing that cellular subfunctionalization arises through the co-option of existing gene regulatory networks. Together, this comparative single-cell analysis of compound eyes bridges microevolutionary transcriptomic divergence with macroevolutionary cellular innovations.

## Introduction

Across the animal kingdom, distinct eye prototypes have evolved, with each undergoing extensive secondary modifications to meet the demands of diverse visually guided behaviors.^1^ A fundamental question in evolutionary biology remains: how do these complex systems, often constrained by pleiotropy, undergo rapid modifications during species diversification? While comparative studies of visual systems have traditionally focused on opsin evolution^2,3^ and photoreceptor electrophysiology^4,5^, this emphasis has obscured a more comprehensive view of eye evolution and the regulatory mechanisms that drive visual novelties. The advent of single-cell (scRNA-seq) and single-nucleus RNA sequencing (snRNA-seq) have enabled the systematic molecular characterization of cell types, facilitating robust comparative analyses of cell type evolution. However, existing comparative eye atlases are predominantly restricted to the camera-type eyes of vertebrates^6,7^ and cephalopods^8,9^. While the visual system of *Drosophila melanogaster* has been comprehensively characterized at the single-cell level,^10–12^ the evolutionary dynamics of cell types in the compound eye, the other major eye type, remain poorly understood. Furthermore, most comparative eye atlases focus on distantly related taxa, hindering the understanding of the origins of visual system cell type diversity.

Neotropical *Heliconius* butterflies represent a powerful system for investigating compound eye cell type evolution across short evolutionary timescales. As a classic example of adaptive radiation,^13^ this genus is characterized by the rapid diversification of aposematic wing color patterns within the last 10 million years.^14^ Accompanying this morphological radiation, *Heliconius* evolved a vast diversity of visual behaviors, including visual mate choice based on wing coloration^15–19^ and sexually dimorphic color vision^20,21^. *Heliconius* rely heavily on vision for foraging^22^, host-plant selection^23,24^, and mate recognition^25^, and this heavy reliance on vision is also mirrored by increased visual investment in neuroanatomical structures. Compared to outgroup species in the Heliconiini tribe, the genus *Heliconius* displays a marked expansion of the visual processing areas within the mushroom bodies of the central brain.^26^ Furthermore, substantial variation in compound eye size exists both between species and between sexes.^27,28^ Beyond these neuroanatomical shifts, opsin sensitivity, expression, and the resulting spectral sensitivity of photoreceptors also exhibit significant diversity.^21,29–35^ However, aside from the stochastic expression of *spineless* (*ss*), which determines the pale and yellow ommatidial types,^36,37^ the regulatory mechanisms that partition these diverse butterfly opsins into distinct photoreceptors remain largely unknown. Moreover, while a few recent studies have begun to connect genetic variation to visual behaviors,^20,38,39^ comparative analysis connecting these genetic variants to specific cell type changes, and ultimately to behavioral divergence, remain scarce.

In this study, we present a comparative, single-cell resolution analysis of compound eye evolution by integrating single-nucleus transcriptomics and epigenomics across five *Heliconius* species. By examining taxa ranging from a single population polymorphic for visual mate preference to deeply diverged species pairs, we uncover the multi-modal changes driving sensory diversification. To link these molecular innovations to visual behavior, we utilize population genomics and chromatin conformation capture assays. In polymorphic *H. cydno alithea*, this approach identifies *senseless-2* (*sens-2*) as the primary visual preference gene. Unexpectedly, we find that *sens-2* is expressed not in neurons, but in a specialized fenestrated glial cell type. In contrast, across species with fixed mate preferences (*H. cydno galanthus*, *H. pachinus*, and *H. melpomene*), photoreceptors emerge as the most divergent cell type, exhibiting differential expression of mate-preference genes. We demonstrate that accelerated transcriptomic divergence in photoreceptors is a general pattern, leading to the evolution of novel photoreceptor subtypes and compositional shifts in these subtypes. Together, these findings illustrate how distinct cellular and regulatory mechanisms, spanning diverse nervous system cell types, shape the evolution of the compound eye at different stages of divergence.

## Results

### De novo genome assembly reveals a complex repeat cluster at the Heliconius cydno mate preference locus

The polymorphic subspecies *Heliconius cydno alithea* in western Ecuador provides a unique opportunity to study compound eye diversification in the earliest stage of speciation.^40^ In this subspecies, white and yellow morph individuals co-occur throughout much of its range, and this wing coloration difference is associated with divergent male visual mate preference: yellow males, on average, show preference for yellow females, whereas white males show no preference as a group.^17,41^ We previously conducted a GWAS of male mate preference in *H. c. alithea*,^34^ and found significant associations in a roughly 1 Mb region on chromosome 1, also known as the *K* locus. In addition to the putative preference genes, this locus also contains *aristaless-1* (*al-1*), which regulates the white-versus-yellow forewing color switch.^42^ However, the causal genetic variants underlying mate preference variation remained unclear due to a repeat-rich unassembled gap in the center of the *K* locus.

To fully resolve this locus, we generated a *de novo*, chromosome-level genome assembly of *H. c. alithea* using Oxford Nanopore long-reads (∼80x coverage; read N50 = 23.2 Kb) and Micro-C chromatin conformation capture data. This resulted in a highly-contiguous assembly (contig N50 = 14.9 Mb) of all autosomes and the Z chromosome with only 11 gaps (Fig. S1). The assembly had a high gene completeness (99.0%) using BUSCO Lepidoptera single-copy orthologues^43^ (odb10; n = 5286). We found that the previous *K* locus gap comprised a 65 kb repeat-rich 5S ribosomal DNA cluster. Long-read sequencing from additional individuals and species revealed that this region was highly variable in size and exhibited a complex history of rearrangements within the *cydno* clade (Fig. S2).

### The polygenic architecture of mate preference in H. c. alithea

The *Heliconius* genome exhibits a large amount of structural variation (SV).^44^ To resolve genetic variants potentially missed by our previous reference-based approaches, we implemented a dual, complementary GWAS strategy: a standard SNP-based analysis utilizing our new chromosome-level reference, and a reference-free *k*-mer association study. Following the kmerGWAS pipeline^45^, this reference-free approach utilizes short sequences of fixed length (*k*-mers) derived from sequencing data to identify the presence or absence of SVs, such as insertions and deletions. We performed GWAS for the 1,481 previously recorded mate choice events made by 109 males,^34,41^ using a generalized linear mixed model implemented in the GMMAT^46^ to test associations independently for 7 million SNPs and the presence/absence of 126 million 25-mers. Using a FDR threshold of 0.01 derived from our SNP-based GWAS (p < 1.86e-08), we identified 225 significant *k*-mers (Table S1), many of which capture variants from alternative haplotypes missing in our reference genome. To identify the genomic location of these variants, we further assembled reads containing significant *k*-mers and mapped the consensus contigs to our reference.

Our *k*-mer approach was supported by the recovery of the most significant SNP at 15.45 Mb (Fig. 1B). Notably, the most significant *k*-mer mapped to a non-repetitive region within the 5S rDNA cluster (Fig. 1B). In the GWAS population, the absence of this *k*-mer shifted average male preference for white females from 63% to 37% (*p* < 5×10^-11^, Fig. 1C). Furthermore, we identified additional association peaks on chromosome 11 and 17 (Fig. 1A; Fig. S3). Intriguingly, the effect sizes for significant *k*-mers on chromosome 1 were consistently positive whereas those on chromosome 11 were negative, suggesting distinct mechanisms controlling white or yellow preference. This modular genetic architecture mirrors the distinct QTLs on separate chromosomes that govern red and white preference in *H. melpomene* and *H. cydno* hybrids.^47^

**Figure 1:**
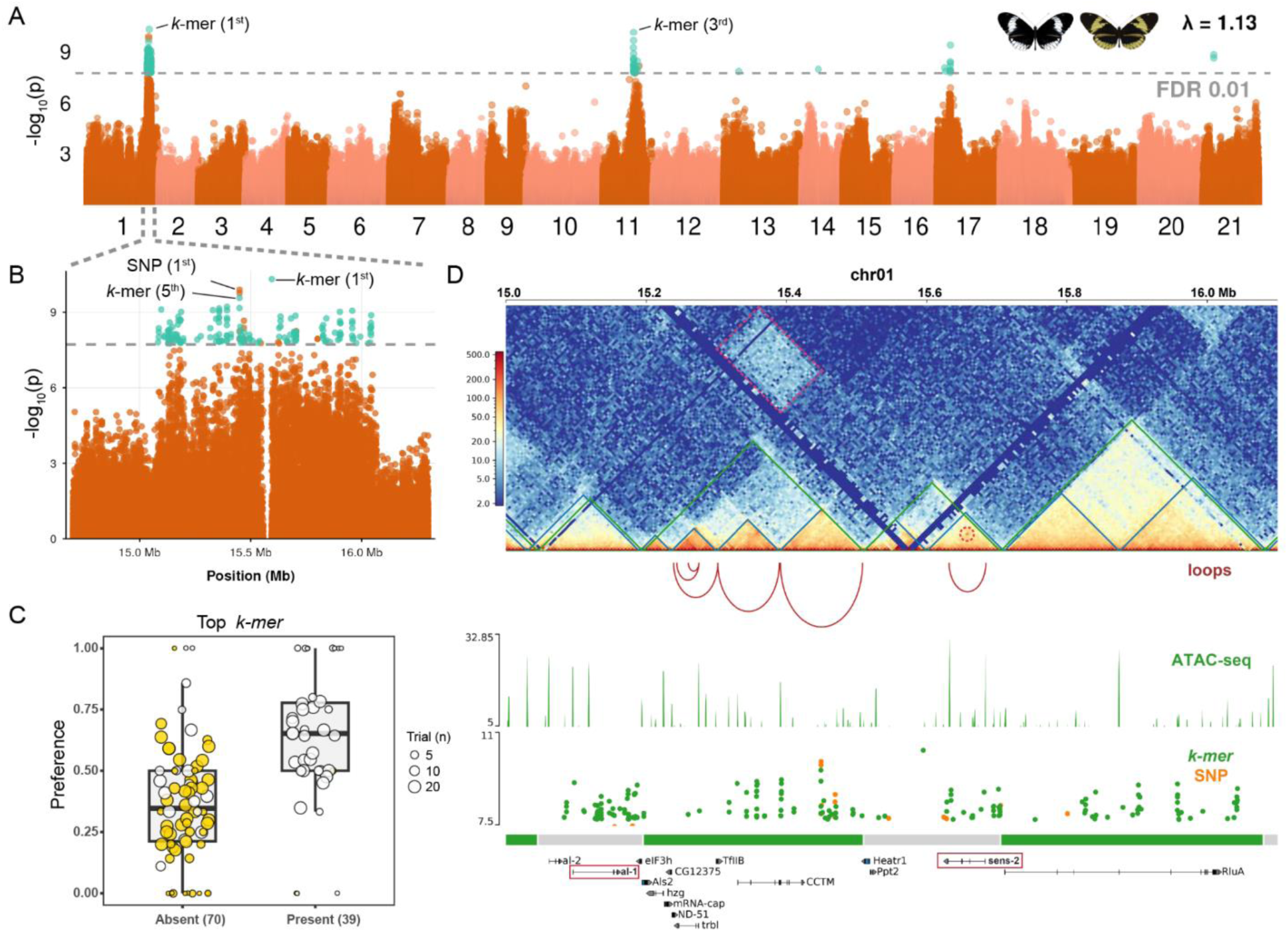
GWAS and 3D chromatin architecture at the male mate preference locus. **(A)** Manhattan plot of the genome-wide association study (GWAS) for male mate preference. Orange and cyan points represent SNP– and *k*-mer-based associations, respectively. Only *k*-mers passing a false discovery rate (FDR) < 0.01 are displayed. **(B)** Zoom-in view of the Manhattan plot across the 2 Mb region on chromosome 1, showing clusters of significantly associated *k*-mers. **(C)** Boxplot illustrating group-level male mate preference based on the absence or presence of the top-associated *k*-mer. Points represent individual males (colored by white or yellow wing coloration), with dot size scaled to the number of behavioral trials (n). **(D)** Integration of 3D chromatin topology and functional genomics at the *K* locus. *Top:* KR-normalized Micro-C contact frequency matrix (5 kb resolution) from adult brain tissue. Green and blue triangles demarcate topologically associating domains (TADs; 15 kb resolution) and sub-TADs (5 kb resolution), respectively. Significant chromatin looping interactions are denoted by red arcs. *Middle:* Chromatin accessibility (ATAC-seq) track from pupal optic lobe tissue. *Bottom:* Fine-scale GWAS signals (orange = SNPs, green = *k*-mers) aligned with TAD boundaries and gene models, highlighting *al-1* and *sens-2*.

In the *K* locus region, we found multiple GWAS association peaks in both SNP-based and *k*-mer-based GWAS, spanning from 15 Mb to 16 Mb, similar to our previous GWAS^34^ (Fig. 1B). A broad elevation of association like this could be indicative of locally suppressed recombination, perhaps due to chromosomal inversions.^48^ However, we found no evidence of such structural rearrangements spanning the *K* locus. Principal component analysis (PCA) of SNPs in our GWAS population using Asaph^49^ revealed no inversions (Fig. S4), and direct structural variant calling from long reads via SVIM^50^ confirmed the absence of inversions. This aligns with recombination maps in *H. melpomene* and *H. cydno*, which similarly show no evidence of megabase scale inversions.^51^ Furthermore, Linkage Disequilibrium (LD) decay analysis demonstrated a rapid decline to background levels outside the association peaks in the *K* locus.^34^ Collectively, these results suggest that the broad association signal is not due to repressed recombination, but rather the result of selection acting on multiple distinct loci within the *K* locus.

### Long-range interactions between color and preference loci

To further investigate the mechanism driving this GWAS association pattern and prioritize candidate genes for mate preference, we generated high-resolution Micro-C maps from the whole brains of adult white and yellow *H. c. alithea* males. Enhancer-promoter interactions are largely constrained within topologically associating domains (TADs).^52,53^ We identified four TADs and nine sub-TADs, at 20-kb and 5-kb resolution, respectively, in the *K* locus (Fig. 1D). Notably, we found a highly conserved TAD organization between white and yellow males (Fig. S5).

Within the *K* locus, the TAD containing the forewing color-switch gene *al-1* exhibited high contact frequency with another TAD 600 kb away (Fig. 1D), which contained the most significant mate preference GWAS *k*-mer and the protein-coding gene *sens-2*. *sens-2* encodes a transcription factor with six C2H2 zinc finger (ZF) domains, a distant paralog of four-ZF Senseless (Sens).^54^ While *sens* is well-known for its role in photoreceptor differentiation,^55,56^ the function of *sens-2* remains largely uncharacterized, having previously only been reported in the *Drosophila* embryonic midgut^57,58^ and larval surface glia^59^. The A/B compartment analysis further revealed that these two interacting TADs are within the same B compartment (Fig. S6), typically associated with transcriptionally repressed or heterochromatic states.

Similarly, an adjacent TAD situated just upstream of the *K* locus was also in the B compartment (Fig. S6). This TAD contains *neither inactivation nor afterpotential C* (*ninaC*), a highly expressed photoreceptor marker gene.^60^ Robust inter-TAD long-range interactions suggest a coordinated regulatory mechanism, where the wing color gene *al-1*, the candidate mate preference gene *sens-2*, and a key phototransduction gene *ninaC* are coupled in 3D chromatin architecture, which can facilitate synchronized transcriptional activation or silencing.

To map fine-scale regulatory interactions within the *K* locus, we integrated Micro-C chromatin loops with chromatin accessibility profiles we generated from adult brains via ATAC-seq^61^. While most loop anchors coincided with structural TAD boundaries, we identified a distinct intra-domain loop linking the *sens-2* promoter to a putative intronic enhancer (Fig. 1D). Both loop anchors exhibit high chromatin accessibility, supporting a functional regulatory contact. The maintenance of this loop in the adult brain, despite low expression of *sens-2* (Fig. S7), is consistent with the *Drosophila* model where enhancer-promoter topologies are established prior to transcriptional activation^62^. In summary, Micro-C data revealed a multi-scale 3D DNA architecture at the mate preference locus, characterized by long-range inter-TAD contacts linking the wing color gene *al-1* and the candidate preference gene *sens-2*, coupled with fine-scale enhancer-promoter looping at *sens-2*.

### Compound eye cell atlas of H. c. alithea

While Micro-C linked GWAS variants to candidate genes within specific TADs, the cellular mechanisms driving this visual mate preference remained unresolved. The butterfly visual system consists of the retina, optic lobe, and central brain. The retina or compound eye is built from repeating ommatidia, each consisting of photoreceptors and retinal glia, including cone and pigment cells^63^ (Fig. 2A). While each *Drosophila* ommatidium contains eight photoreceptors, with the R7 cell subdivided into pale (pR7) or yellow (yR7) subtypes based on opsin expression^37^, most butterflies possess nine photoreceptors per ommatidium, including two R7 homologs^36^. Visual information perceived by photoreceptors is transmitted directly to the underlying lamina and medulla of the optic lobe, which contains its own neurons and glia^64^.

**Figure 2:**
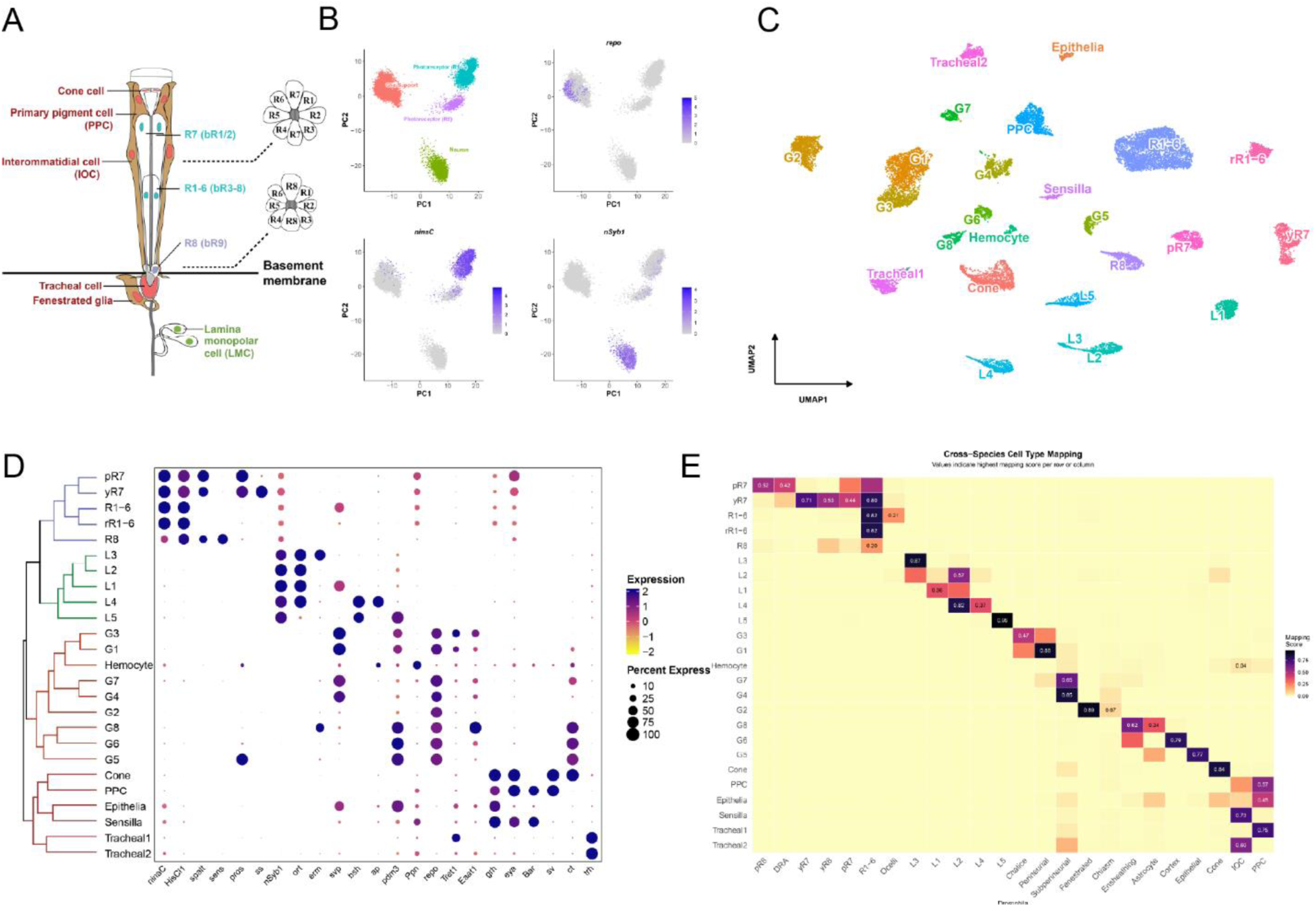
A single-nucleus transcriptomic atlas of the *Heliconius* compound eye. **(A)** Schematic anatomy of a butterfly ommatidium. Nuclei are color-coded by major cellular lineages: support cells (red) and photoreceptors (blue and purple). Photoreceptor nomenclature reflects *Drosophila* homology, with traditional butterfly terminology (b-prefix) provided in parentheses. **(B)** Principal component analysis (PCA) of 20,586 nuclei from the adult *H. c. alithea* compound eye (left), alongside feature plots displaying the expression of canonical lineage markers: *repo* (glial cells), *ninaC* (photoreceptors), and *nSyb1* (non-photoreceptor neurons). **(C)** Uniform Manifold Approximation and Projection (UMAP) embedding of the integrated snRNA-seq dataset, colored and annotated by cell-type cluster at Leiden resolution of 0.5. **(D)** Expression dotplot of marker genes across the identified cell types. Dot size indicates the percentage of nuclei expressing the gene, and color intensity represents the scaled average expression. The dendrogram illustrates hierarchical clustering of cell types based on the transcriptomic similarity of highly variable genes within the Harmony embedding space. **(E)** Cross-species transcriptome alignment heatmap mapping *Heliconius* cell types (rows) to *Drosophila* cell types (columns) derived from the curated eye Fly Cell Atlas. Color intensity reflects the mapping score.

To identify the specific cell types expressing *sens-2* and other candidate genes, we integrated our GWAS and chromatin architecture results with single-cell transcriptomics. We generated single-nucleus RNA-seq (snRNA-seq) data from the compound eyes (retina and lamina) of 10 adult male *H. c. alithea* (Table S2-S3). Gene quantification was performed using a custom *de novo H. c. alithea* annotation with refined untranslated regions (UTRs) to maximize transcriptome read recovery. Following stringent quality control, including doublet removal and ambient RNA correction, we recovered a total of 20,586 high-quality nuclei.

For initial broad cell type group assignments, we performed PCA on the 979 highly variable genes identified by the S–E model implemented in ROGUE^65^. The first two PCs revealed three major groups: PC1 partitioned the nervous system into neuronal lineages, marked by *neuronal Synaptobrevin* (*nSyb*)^66^, and glial lineages, defined by *reversed polarity* (*repo*)^67^; while PC2 further distinguished photoreceptors, marked by *ninaC*, from downstream optic lobe neurons (Fig. 2B). To resolve fine-scale cell types, we performed Leiden clustering (resolution = 0.5) on the Harmony^68^-integrated dataset, which corrected the individual batch effects. This unsupervised approach identified 25 distinct clusters (Fig. 2C). We hierarchically organized these clusters into a dendrogram based on their transcriptomic similarities of the Harmony embedding. Cell type identities were assigned using established *Drosophila* or *Papilio* butterfly marker genes for photoreceptors^36,69,70^, lamina neurons^70,71^, retinal glia^72,73^, and tracheal cells^74^ (Fig. 2D).

To complement our marker-based annotations and resolve glial populations, which typically lack canonical markers, we mapped our *Heliconius* atlas to the *Drosophila* Fly Cell Atlas (FCA)^12^ using SAMap^75^. However, initial direct alignment yielded ambiguous results. To resolve these inconsistencies, we implemented a supervised refinement approach: we mapped three *Drosophila* scRNA-seq datasets from the adult retina^10^, optic lobe^11^, and glia^76^ onto the subsetted FCA whole head atlas using scANVI^77^. This allowed us to update inconsistent FCA cell type annotations, from which we derived a high-quality *Drosophila* eye snRNA-seq reference. Using SAMap to transfer labels from the curated *Drosophila* reference to the *Heliconius* atlas, we were able to map 23 of the 25 *Heliconius* clusters to *Drosophila* clusters (mapping score > 0.3), despite 300 million years of divergence between Lepidoptera and Diptera^78^.

Overall, the cell-type labels predicted by SAMap were consistent with our marker-based annotations (Fig. 2E). However, the *Heliconius* R8 photoreceptor, which was marked by *sens*^36,56^, lacked a strong mapping to any *Drosophila* cell type. This transcriptomic divergence likely reflects its specialized anatomical structure, as the butterfly R8 is basally restricted and contributes only a minor fraction to the rhabdom. Furthermore, the *Drosophila* primary pigment cells (PPCs) and interommatidial cells (IOCs) exhibited ambiguous, many-to-one mappings from multiple butterfly support cell types. Our atlas also uncovered compound eye cell types unique to *Heliconius*. For instance, we identified two distinct tracheal cell types not found in the *Drosophila* head FCA, both expressing the tracheal cell fate gene *trachealess* (*trh*)^74^. These likely represent the cellular components of the tapetum, the specialized reflective terminal tracheae characteristic of Lepidoptera^79^.

### Glial expression of the mate preference gene sens-2

Leveraging our newly established *Heliconius* eye atlas, we sought to identify the cell types associated with mate preference variation by investigating the expression profiles of GWAS candidate genes. To identify causal behavioral genes, which are expected to show cell-type– or individual-specific expression patterns, we calculated expression entropy (S) and maximal S-reduction (*ds*) using the S–E model implemented in ROGUE^65^.

We identified 98 genes sharing a sub-TAD with significant *k*-mers. Of these, four candidates showed significant *ds* scores (adjusted *p* < 0.05): *sens-2* and *RluA pseudouridine synthase* (*RluA*) from the *K* locus, sidestep VI (*side-VI*) from chromosome 17, and *CG43373* (predicted to encode adenylate cyclase) from chromosome 21 (Fig. 3A). When mapped to *Heliconius* cell clusters, *RluA* and *CG43373* showed broad expression across subsets of both neuronal and glial populations. In contrast, *side-VI* expression was restricted to pale R7 (pR7) photoreceptors and L4 neurons (Fig. 3B), while *sens-2* was highly enriched in the G2 glial cell type (Fig. 3C), which maps to *Drosophila* fenestrated glia (Fig. 2E). *In situ* hybridization using hybridization chain reaction (HCR) on adult *H. c. alithea* compound eye sections corroborated our snRNA-seq findings. Expression of *sens-2* was localized immediately beneath the basement membrane of the retina, corresponding to the fenestrated layer, where bundled photoreceptor axons pass through (Fig. 3D). This retina-specific expression pattern of *sens-2* was further supported by bulk RNA-seq data, which demonstrated that *sens-2* is consistently enriched in the retina relative to the optic lobe and central brain throughout pupal development (Fig. S7).

**Figure 3:**
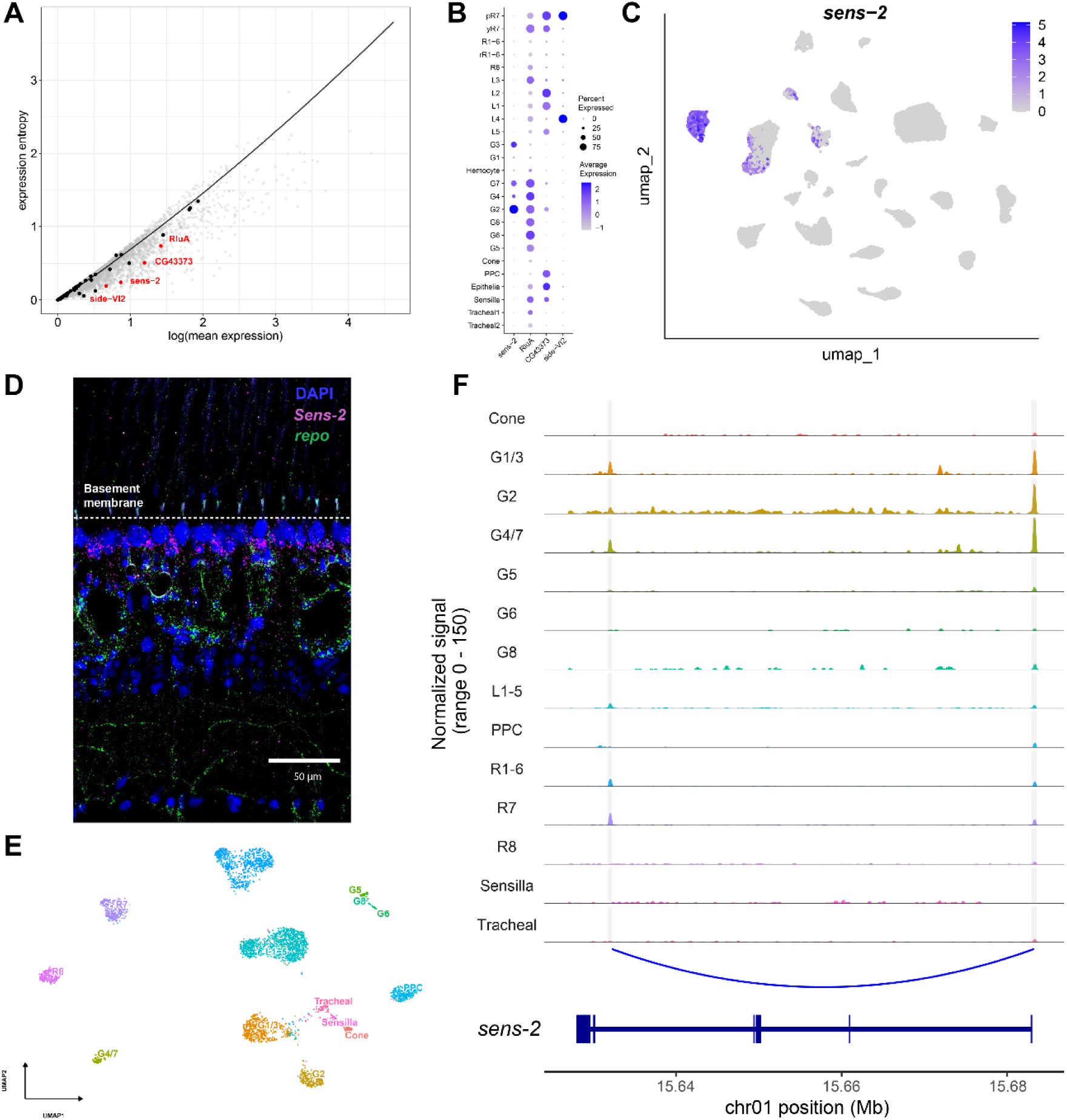
Cell-type-specific expression and cis-regulatory landscape of *sens-2*. **(A)** S–E plot illustrating expression entropy (S) versus log mean expression (E) across all detected genes in the *H. c. alithea* snRNA-seq. The solid black line represents the null model LOESS regression fit. Black dots denote candidate genes located within TADs containing significant mate preference GWAS variants. Candidates exhibiting significant maximal S-reduction (*ds*) are highlighted in red and labeled. **(B)** Dot plot of the four significant candidate genes, with dot size representing the percentage of expressing nuclei and color indicating the average expression levels. **(C)** UMAP feature plot of *sens-2* expression. **(D)** HCR staining of the adult *H. c. alithea* retina. *sens-2* (magenta) strongly localized to nuclei within the fenestrated glia cell layer. *repo* was utilized alongside it as a pan-glial marker to define this layer. **(E)** UMAP of adult *H. c. galanthus* snATAC-seq nuclei. Cell type identities were assigned via integration with the snRNA-seq atlas. **(F)** Cell-type-specific chromatin accessibility tracks at the *sens-2* locus. Rows represent aggregated Tn5 transposase insertion frequencies per cell type. Gray boxes highlight the putative enhancer (left) and the promoter (right) of *sens-2*.

To determine whether *sens-2* expression in fenestrated glia is conserved, we compared its localization in *Heliconius* and *Drosophila*. Using Sens-2 antibody staining and FCA single-cell transcriptomic data, we confirmed that *sens-2* expression is similarly restricted to fenestrated glia in the adult *Drosophila* head (Fig. S8). Next, to investigate the potential functions of G2 fenestrated glia in *Heliconius*, we performed gene ontology (GO) enrichment analysis on G2 marker genes. This analysis revealed an overrepresentation of terms related to eye pigmentation and the ommochrome synthesis pathway (Fig. S9, Table S4), suggesting that *Heliconius* fenestrated glia are pigmented, similar to those in *Drosophila*^80^. While we sought to functionally validate the role of *sens-2* in G2 glia by knocking out *sens-2* using CRISPR/Cas9, we recovered extremely few individuals with mutations in the pupal stage (Table S5-S7), a result consistent with the pupal stage lethality reported for *sens-2* mutants in *Drosophila*^59^.

We next investigated the cis-regulatory mechanisms governing the expression of *sens-2* in glial cells. We performed single-nucleus ATAC-seq (snATAC-seq) on the adult *H. cydno* retina to characterize chromatin accessibility across cell types. After rigorous quality control, we recovered 4,140 high-quality nuclei and 79,635 distinct open chromatin regions. To annotate these nuclei, we used Signac^81^ multimodal label transfer to map the snATAC-seq data against our snRNA-seq reference atlas (Fig. 3E). To identify cell-type-specific accessible chromatin regions, we performed differential accessibility analysis comparing each cell type to all others combined. At the *sens-2* gene body, the promoter region was highly accessible in G1/3, G2, and G4/7 cells (Fig. 3F). In contrast, the putative enhancer identified from Micro-C showed stronger accessibility only in R7 photoreceptor and G1/3 glia (Fig. 3F), suggesting the potential role of *sens-2* in photoreceptor development.

In summary, we identified *sens-2* as a key candidate gene for mate preference. Through GWAS and Micro-C, we found that *sens-2* is located within a TAD that physically contacts the color gene and contains the top GWAS *k*-mer. Subsequent snRNA-seq revealed that *sens-2* exhibits highly cell-type-specific expression. Surprisingly, while its paralog, *sens*, is expressed in R8 photoreceptors, *sens-2* is highly expressed in fenestrated glia, a finding we corroborated using *in situ* hybridization. Furthermore, snATAC-seq and Micro-C showed high accessibility of a *sens-2* enhancer that forms a chromatin loop with the *sens*-2 promoter specifically in R7 photoreceptors, suggesting an additional role in photoreceptors. Because glia are known to influence photoreceptors by recycling neurotransmitters^82,83^ and guiding axons^59,84^ during development, we hypothesize that *sens-2* impacts color information processing by altering inter-photoreceptor synapse formation during early development or by modulating synaptic strength through histamine recycling at the fenestrated layer.

### Distinct mechanisms of gene expression and cell type evolution between species pairs

Having characterized genes and cell types associated with mate preference in polymorphic *H. c. alithea*, we next investigated whether these mechanisms are conserved across divergent species pairs. We focused on three lineages exhibiting strong, assortative male mate preference: the parapatric sister species *H. cydno galanthus* (white) and *H. pachinus* (yellow), alongside their respective sympatric populations of the outgroup *H. melpomene* (red) from the Caribbean and Pacific coasts of Costa Rica (Fig. 4A). Hybrids between *H. c. galanthus* and *H. pachinus* are fully fertile^85^, whereas crosses between either species and *H. melpomene* yield sterile heterogametic females^86^. While our previous work identified species-level differences in opsin expression, filtering pigments^35^, and photoreceptor synaptic connections^34^, the cell types associated with mate preference across these species remain unresolved.

**Figure 4:**
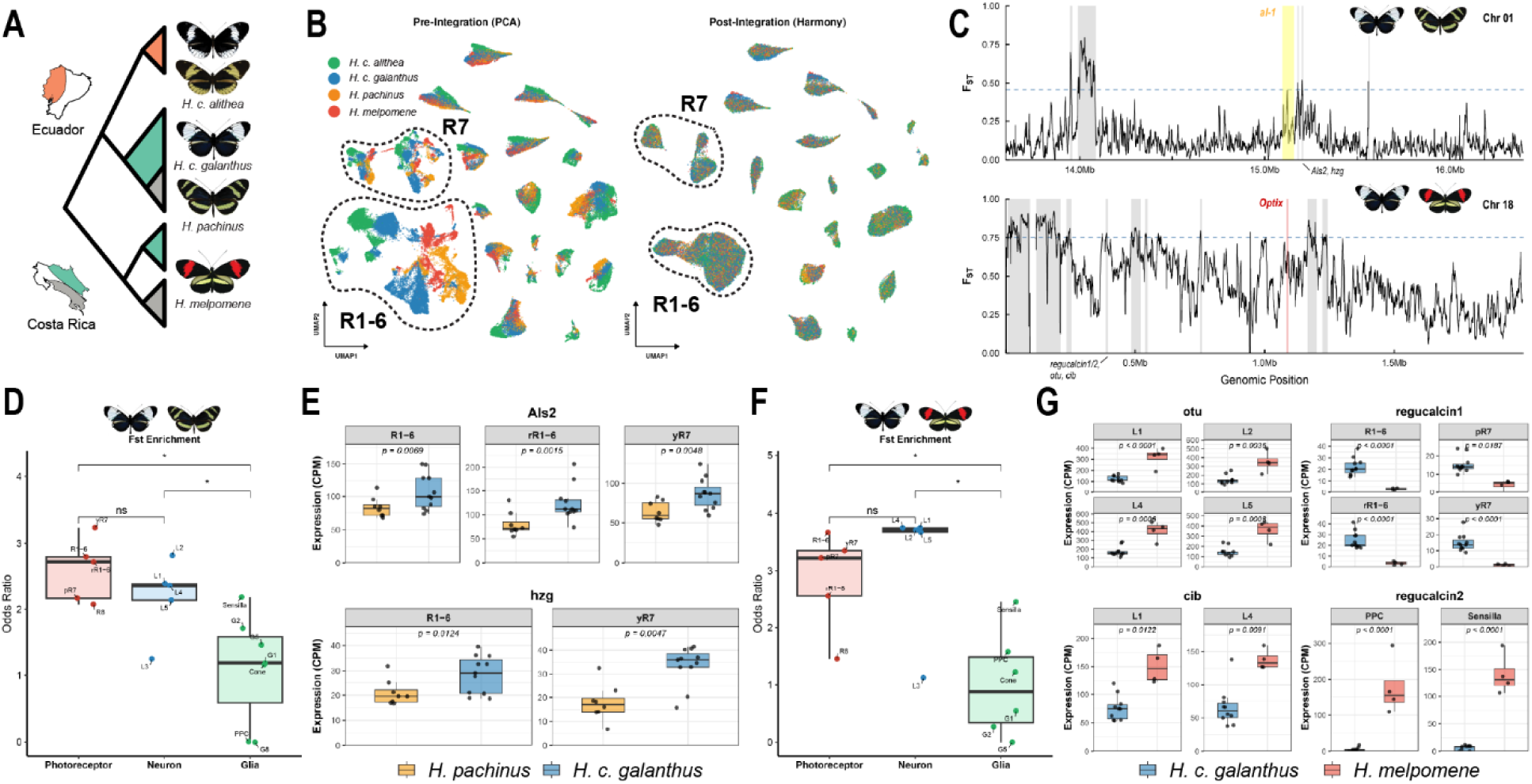
Cellular divergence of mate preference loci across species with divergent mate preference. **(A)** Phylogenetic relationships within the *H. cydno* complex and *H. melpomene*. *H. c. galanthus* (white-preferring) and *H. pachinus* (yellow-preferring) are distributed in Costa Rica. Polymorphic *H. c. alithea* (Ecuador) is an outgroup to the Costa Rican *H. cydno*. *H. melpomene* (red-preferring) is the outgroup to the entire *H. cydno* complex. **(B)** Retina snRNA-seq UMAPs before and after Harmony integration. Each nucleus is colored by species. Dashed lines highlight the R7 and R1-6 photoreceptors. **(C)** Sliding window *F*_ST_ (5 kb) at mate preference loci between pairs of species with divergent male mate preference. *Top:* Comparison between *H. pachinus* and *H. c. galanthus* (Chromosome 1 white-versus-yellow preference QTL) with the top 0.5% *F*_ST_ threshold (dashed line) and the white color gene *al-1* (yellow shading). *Bottom:* Comparison between *H. c. galanthus* and *H. melpomene* (Chromosome 18 white-versus-red preference QTL) with the top 1% *F*_ST_ threshold and the red color gene *optix*. Gray shaded boxes denote *F*_ST_ windows passing the threshold, putative preference genes are labeled. **(D and F)** Enrichment odds ratios for top *F*_ST_ genes among cell-type-specific differentially expressed genes for **(D)** male *H. c. galanthus* vs. *H. pachinus* and **(F)** male *H. c. galanthus* vs. *H. melpomene*. **(E and G)** Cell-type-specific pseudobulk expression boxplots for the candidate preference genes highlighted in panel C. Each dot is the pseudobulk expression of one male individual.

We generated single-nucleus transcriptomes from the eyes of adult males and females for each species and integrated these data with our *H. c. alithea* reference atlas. This cross-species integration revealed a strong conservation of retinal cell types except for the R1-6 and R7 photoreceptors (Fig. 4B). We observed no evidence of cell type gain or loss, suggesting that behavioral divergence at this stage arises from cell abundance differences or transcriptomic divergence of the existing cell types.

To identify the cellular drivers of preference between *H. pachinus* and *H. c. galanthus*, we first focused on the genomic loci associated with mate preference variation. Previous QTL mapping identified a major mate-preference locus on chromosome 1^16^. Using the estimated recombination rate of 3.17 cM/Mb^51^ and *wingless* as a gene marker anchor, we refined this QTL interval to a physical region spanning 13.6–16.4 Mb in the *H. c. alithea* reference genome. Notably, the *K* locus (15-16 Mb) that we identified in *H. c. alithea* was fully nested within this broader interval. We prioritized candidate genes by scanning for regions of high differentiation using *F_st_* within this locus. Using a stringent *F_st_* cutoff (top 0.5% of 5kb genome-wide sliding windows), we identified five peaks of elevated differentiation and three out of five peaks fell within the previously defined *K* locus (Fig. 4C).

We reasoned that the cell types most relevant to this behavioral divergence would be enriched for genes showing both DNA-level divergence (genome-wide *F_st_* outliers) and differential expression (DE). By calculating the odds ratio of this overlap between cell-type specific DE genes and genes within *F_st_* outlier windows, we found that photoreceptors and lamina neurons were highly divergent between *H. pachinus* and *H. c. galanthus*, as they exhibited significantly higher enrichment odds ratios compared to glia (Wilcoxon Rank-Sum Test p < 0.05; Fig. 4D). Within neurons, the yR7 photoreceptor subtype displayed the highest enrichment odds ratio, implying it is the most divergent cell type between *H. pachinus* and *H. c. galanthus* adult eyes. Consistent with the genome-wide pattern of highly divergent photoreceptors, two candidate preference genes within the *K* locus, *Amyotrophic lateral sclerosis 2* (*Als2*) and *herzog* (*hzg*), were localized within *F_st_* outlier windows and differentially expressed between *H. pachinus* and *H. c. galanthus* males in both yR7 and R1-6 photoreceptors (Fig. 4D).

Although *sens-2* is not significantly differentially expressed between *H. pachinus* and *H. c. galanthus* males, one of the three *K* locus *F_st_* outlier windows was located within the *sens-2* TAD, near the 5S rDNA cluster (Fig. 4C). Intriguingly, when comparing two *H. melpomene* populations that exhibit divergent mate preferences (white versus yellow) driven by reinforcement^18^, the maximum *F_st_* window within the *K* locus similarly mapped to the 5S rDNA cluster (Fig. S10). This parallel genomic signature suggests that this specific region acts as a recurrent hotspot for the evolution of behavioral isolation.

Finally, we examined the cellular basis of the species-level divergence in mate preference between white *H. cydno* (white-preferring) and *H. melpomene* (red-preferring). This behavioral isolation is largely governed by a quantitative trait locus (QTL) on chromosome 18 driving red preference^47^. Recent work identified *regucalcin1* from this QTL as the gene underlying the mate preference for red patterns^39^. However, while the previous study focused exclusively on *regucalcin1* expression in the brain^39^, our single-nucleus atlas revealed enrichment of this gene in the periphery, specifically within photoreceptors. Across all photoreceptor subtypes except R8, *H. c. galanthus* males exhibited significantly elevated expression of *regucalcin1* compared to *H. melpomene* males. Intriguingly, its tandem duplicate, *regucalcin2*, demonstrated the inverse pattern, with higher expression in the pigment cells of *H. melpomene* (Fig. 4G). Furthermore, two additional genes, *ovarian tumor* (*otu*) and *ciboulot* (*cib*), located within this top 1% *F_st_* window were significantly upregulated in the lamina neurons of *H. melpomene*. This expression shift in lamina neurons aligns with the genome-wide pattern that lamina neurons display a strong enrichment for *F_st_* outliers among cell-type-specific DE genes (Fig. 4F). Overall, our findings suggest that photoreceptors, particularly the yR7 subtype, drive the divergence in white-versus-yellow preference between *H. c. galanthus* and *H. pachinus*. In contrast, the white-versus-red preference between *H. c. galanthus* and *H. melpomene* appears to be driven by a broader divergence across both lamina neurons and photoreceptors.

### Distinct evolutionary rates across compound eye cell types

The evolution of the compound eye is largely driven by selection on diverse visually guided behaviors^1^. While we have focused thus far on visual mate preference and its associated cell type variation, *Heliconius* species also inhabit distinct microhabitats that require the identification of host plants and food sources under varying light conditions^87,88^. To elucidate the broader evolutionary dynamics of compound eye cell types, we expanded our focus beyond preference-associated genes to investigate transcriptome-wide divergence and compositional shifts.

To establish a phylogenetic tree across populations for this comparative analysis, we performed whole-genome sequencing of *H. c. alithea*, *H. pachinus*, *H. c. galanthus*, and the outgroup *H. melpomene* used in snRNA-seq. Phylogenomic reconstruction using a concatenated SNP matrix recovered the sister relationship between polymorphic *H. c. alithea* (Ecuador) and the Costa Rican clade (*H. pachinus* and *H. c. galanthus*). Notably, the tree topology revealed that *H. c. galanthus* is paraphyletic, with yellow *H. pachinus* nested deeply within the *galanthus* clade (Fig. S11). The divergence time estimate between the sister species *H. pachinus* and *H. c. galanthus* was comparable to the divergence between their respective sympatric *H. melpomene* populations (Pacific and Caribbean), which lack morphological differences (Fig. S11). This pattern indicates that overall genomic divergence in these lineages is primarily driven by geographic isolation.

Previous efforts to quantify cell-type-specific transcriptomic divergence have largely relied on pairwise comparisons, such as Spearman rank correlations between two sister species^89,90^. To explicitly model the cell-type-specific evolutionary rates of gene expression in the eye, we expanded our analysis to encompass six distinct populations (Fig. S11). We generated pseudobulk data by aggregating gene counts by cell type for each individual, enabling us to model gene expression evolution across the broader phylogeny using the Brownian motion-based gene expression evolution model (CAGEE^91^).

We first compared a uniform single-rate model against a two-rate model that partitioned neurons (including photoreceptors) from glia and other support cells. The two-rate model provided a better fit, revealing that neurons evolve at a substantially higher rate (*σ*^2^ = 0.0106) than glia (*σ*^2^ = 0.0005) (Table 1). Subsetting the neuronal lineage further, a two-rate model separating photoreceptors from downstream lamina neurons outperformed a single-rate neuronal model. Photoreceptors exhibited a highly accelerated evolutionary rate (*σ*^2^ = 0.0173) compared to lamina neurons (*σ*^2^ = 0.0013) (Table 1), supporting the hypothesis that peripheral, terminal sensory receptors experience less constraint. Finally, we characterized rate heterogeneity within the photoreceptor classes. Unlike the *Drosophila* R8, the miniature *Heliconius* R8 homolog (bR9) is anatomically restricted to the basal ommatidium (Fig. 2A) and transcriptomically distinct (Fig. 2B). To test if this divergence extended to its evolutionary rate, we compared a baseline one-rate photoreceptor model against a two-rate model (R1-7 vs. R8) and a three-rate model (R1-6 vs. R7 vs. R8). The three-rate model yielded the best overall fit. Under the three-rate model, R8 emerged as the most evolutionarily conserved photoreceptor class (*σ*^2^ = 0.0027), while R7 exhibited the most rapid divergence (*σ*^2^ = 0.0186), evolving at almost twice the rate of the R1-6 class (*σ*^2^ = 0.0090).

**Table 1.** Evolutionary rate parameters estimated from the pseudobulk data.

| Model | Rates (n) | $-\ln L$ | $\sigma^2_1$ | $\sigma^2_2$ | $\sigma^2_3$ |
| --- | --- | --- | --- | --- | --- |
| Full |  |  |  |  |  |
| Full | 1 | 3,278,860 | 0.0051 |  |  |
| **Neuron vs Glia** | 2 | **3,050,950** | **0.0106** | 0.0005 |  |
| Neuron |  |  |  |  |  |
| Neuron | 1 | 2,057,300 | 0.0061 |  |  |
| **PR vs Lamina** | 2 | **1,957,490** | **0.0173** | 0.0013 |  |
| Photoreceptors |  |  |  |  |  |
| R1-8 | 1 | 1,088,220 | 0.0105 |  |  |
| R1-7 vs R8 | 2 | 1,081,390 | 0.0126 | 0.0027 |  |
| R1-6 vs **R7** vs R8 | 3 | **1,077,290** | 0.0090 | **0.0186** | 0.0027 |

### A novel broad-range R7 photoreceptor and its compositional shifts

Because photoreceptors demonstrated the highest evolutionary rate, we subset and reclustered photoreceptors to identify fine-resolution subtypes. Butterfly R7 cells, central to color vision, are known to encompass multiple subtypes based on opsin profiles and spectral sensitivities. *Heliconius* exhibits even greater complexity, with opsin expression patterns defining up to 15 distinct ommatidial variations^29^. In our integrated dataset, the yR7 population further split into two clusters, yielding three total R7 subtypes (Fig. 5A). Based on their predominant opsin expression profiles, we designated these three R7 subtypes as UV, blue, and broad-range (BR) R7s (Fig. 5C). While blue and UV R7 cells have also been found in Papilionidae species, an early-diverging lineage in the butterfly superfamily, the novel co-expression of LWRh and BRh in BR R7s likely represents a unique nymphalid innovation that enables robust color discrimination in the red wavelength range^21,29,33,63,92,93^.

**Figure 5:**
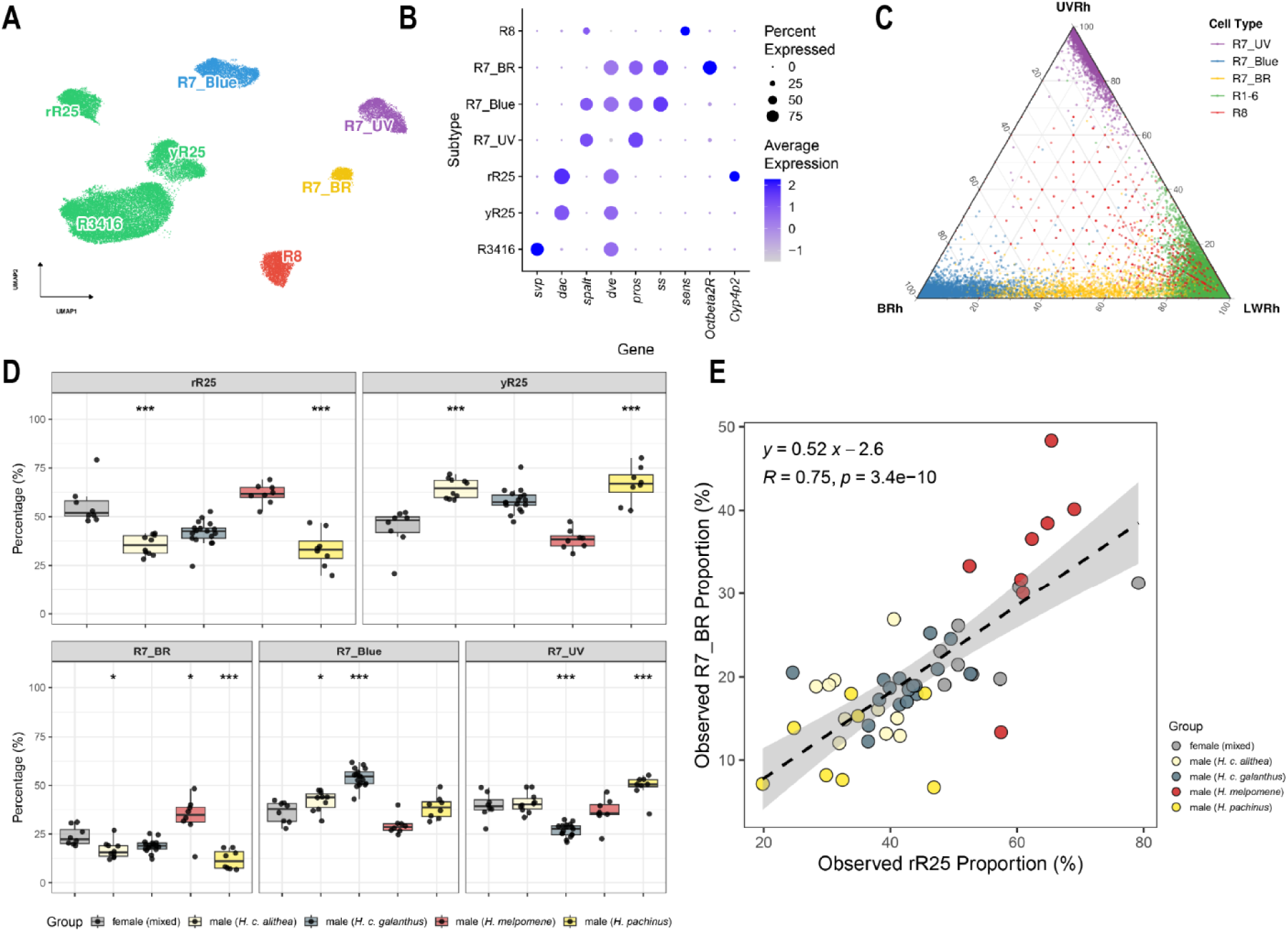
Photoreceptor subtype diversity and proportional shifts across species and sexes. **(A)** UMAP embedding of reclustered photoreceptor nuclei from all species and sex combinations. The R7 lineage segregates into three distinct clusters: UV R7, blue R7, and broad-range (BR) R7. The R1-6 lineage separates into diagonal cells (R3/4 and R1/6) and horizontal cells (R2/5). The R2/5 population further subdivides into two distinct subtypes. **(B)** Dot plot of marker gene expression defining the identified photoreceptor subtypes. **(C)** Ternary plot illustrating relative opsin expression per cell across the photoreceptor populations. **(D)** Relative proportions of the two R2/5 subtypes and the three R7 subtypes across males and females of the four evaluated species. Female samples across all species are aggregated into a single reference group. Asterisks denote significant differences in subtype abundance relative to the aggregated female group. Each data point represents an individual. **(E)** Scatter plot demonstrating the positive correlation between the proportion of the rR2/5 subtype and the BR R7 subtype. Individuals are colored by group.

How did this novel broad-range R7 cell type originate? Cellular novelty often arises through the rewiring of terminal selectors, which are transcription factors that both initiate terminal differentiation during development and continuously maintain neuronal identity in the adult^94^. Therefore, to uncover the terminal selectors defining the BR R7 lineage, we conducted a differential expression (DE) analysis comparing the three mature R7 populations. We found that both BR and blue R7s expressed *spineless*, establishing their homology to the *Drosophila* yellow R7 lineage^37^. In contrast, *Heliconius* UV R7s lacked *spineless*, mirroring the *Drosophila* pale R7^37^ and *Papilio* UV R7^36^. Differential expression analysis further revealed that within the *spineless*-positive photoreceptors, BR R7 lacked expression of the transcription factor *spalt*, which was expressed in other R7 subtypes and R8 cells (Fig. 5B). The stochastic expression of *spalt* in R7 is highly unusual, given its broad conservation in R7 and R8 photoreceptors across developing insect retinas^69^. Loss of the *spalt* complex in *Drosophila* causes R7 and R8 photoreceptors to express R1-6 opsins^95^, a shift that mirrors the expression of LWRh (typically an R1-6 opsin) in BR R7 cells.

Unexpectedly, our subclustering analysis also resolved the broad R1-6 lineage into three distinct populations (Figure 5A). Based on the expression of the marker gene *seven-up* (*svp*), we identified the largest cluster as diagonal photoreceptors (R1/6 and R3/4)^96^. The remaining two clusters represent a previously uncharacterized subdivision of the horizontal photoreceptor class (R2/5). Differential expression analysis of the two subtypes identified the lipid pathway gene *cytochrome P450 4p2* (*Cyp4p2*) as the top marker (Fig. 5B).

We next investigated the compositional changes of the R7 and R2/5 subtypes between males of four lineages and females using scCODA^97^. Because female subtype abundances were highly similar across lineages, we aggregated them into a single reference group for the comparisons. *H. melpomene*, a species inhabiting forest-edge environments, exhibited a significantly higher abundance of BR R7 cells in males (Fig. 5D). In contrast, males of the closed-forest sister species *H. pachinus* and *H. c. galanthus* showed divergent R7 ratios despite their shared habitat. The yellow-preferring *H. pachinus* males displayed an increased proportion of the UV R7 subtype, potentially as an adaptation to detect UV-reflection from yellow wings. Conversely, the white-preferring *H. c. galanthus* males were dominated by the blue R7 subtype. Furthermore, the proportion of BR R7 cells strongly correlated with the proportion of rR2/5 cells (*R* = 0.75) across individuals (Fig. 5E). On average, one BR R7 cell appears for every two rR2/5 cells. This aligns with the observation that BR R7 cells typically appear in ommatidia with red filtering pigments^92^. Thus, the subdivision of R2/5 into two subtypes likely corresponds to red versus yellow ommatidia.

### Cell type subfunctionalization through redeployment of doublesex

Finally, we extended our evolutionary comparison beyond the speciation continuum to investigate the deep divergence between two *Heliconius* subgenera, the *melpomene/cydno* clade and the *erato/sapho* clade (Fig. 6A). These two lineages are separated by 9.6 million years^14^ and exhibit distinct behaviors such as larval feeding behaviors and mating strategies. Most notably, only females in the *erato*/*sapho* clade possess true UV color vision, allowing them to discriminate wavelengths within the UV range. In contrast, males of this clade and both sexes of the *melpomene/cydno* clade cannot^20,21^. This trait is hypothesized to be shaped by both natural selection, enabling females to identify pollen sources, and sexual selection, allowing males to detect the *Heliconius*-specific yellow 3-OHK wing pigment^20^. The ability to discriminate wavelengths in the UV range requires two distinct UV photoreceptor cell types, which rely on the duplication of the ancestral UV opsin gene into *UVRh1* and *UVRh2* at the root of *Heliconius*^30^. While the autosomal location of *UVRh2* is conserved throughout *Heliconius*, *UVRh1* is autosomal in the *melpomene/cydno* clade but resides on the female-specific W chromosome in the *erato/sapho* clade^21^. Although this translocation explains the absence of true UV color vision in the *erato/sapho* clade males, a fundamental regulatory puzzle still remains: How do females subfunctionalize these two UV opsins into distinct photoreceptor populations to achieve true UV vision?

**Figure 6:**
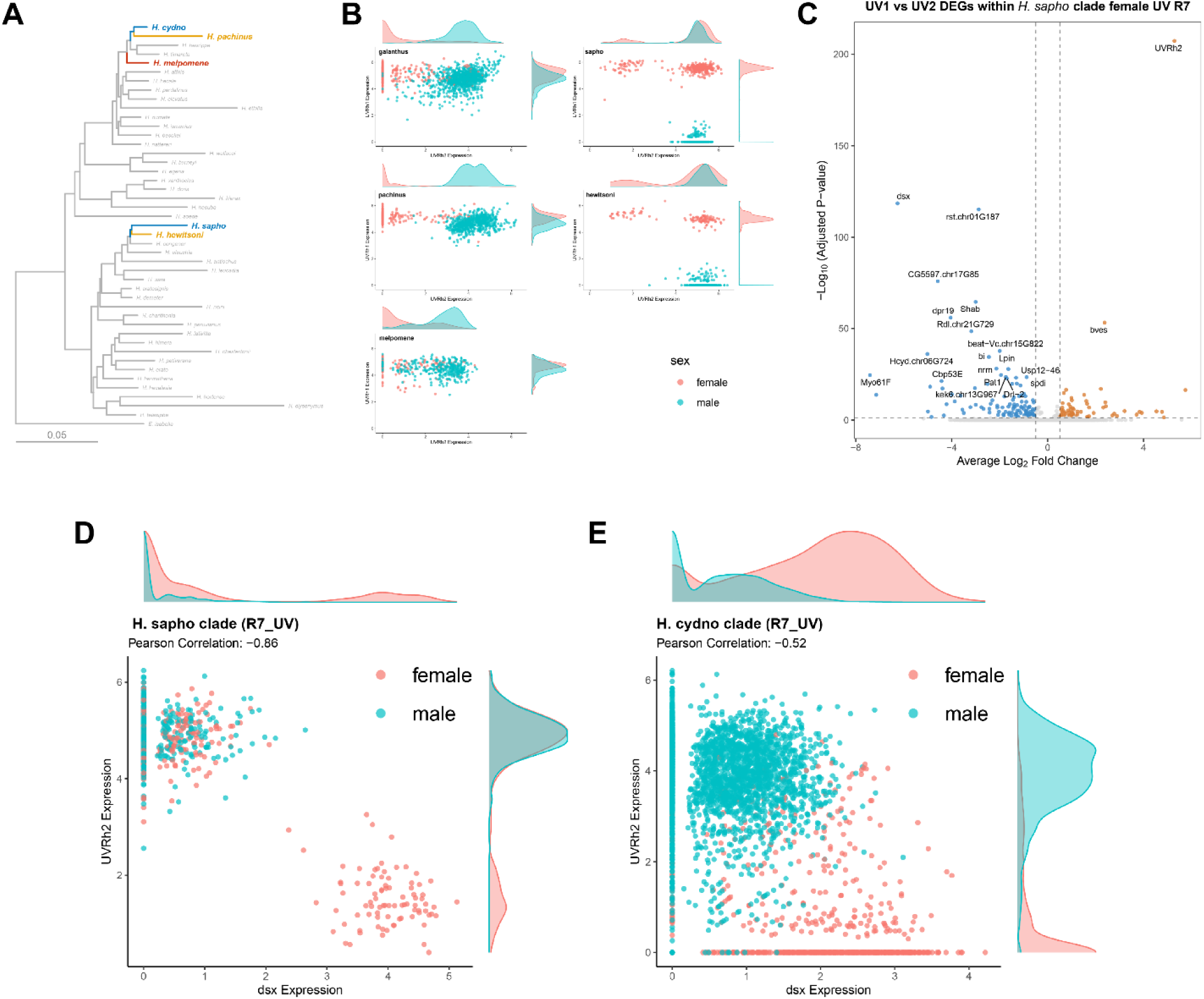
Evolution of a novel female-specific UV R7 subtype in the H. sapho subgenus. **(A)** Phylogenetic tree{Citation} of the *Heliconius* genus. Focal species are bolded and colored by their primary wing coloration (blue, yellow, or red). The *H. sapho* and *H. hewitsoni* lineages belong to a distinct subgenus relative to the *H. melpomene* and *H. cydno* clades. **(B)** Co-expression analysis of *UVRh1* and *UVRh2* in UV R7 photoreceptors across five species, each separated by sex. Density plots reveal a bimodal distribution of *UVRh2* expression specifically within females of the *H. sapho* clade, defining two UV R7 subtypes. **(C)** Differential expression analysis comparing the UV2 (high *UVRh2*) and UV1 (low *UVRh2*) subtypes in *H. sapho* clade females. Top 20 significant marker genes are annotated. **(D and E)** Co-expression scatter plots showing the negative correlation between *dsx* and *UVRh2* within UV R7 cells in **(D)** the *H. sapho* clade and **(E)** the *H. cydno* clade. Cells and density plots are colored by sex.

To resolve the regulatory logic underlying UV cell type subfunctionalization, we generated snRNA-seq profiles from the retinas of male and female *H. sapho* and *H. hewitsoni*, the co-mimics of white *H. c. galanthus* and yellow *H. pachinus,* respectively (Fig. 6A). Integration of these data with our *melpomene/cydno* atlas revealed strong cell type conservation. While males of all species and *melpomene/cydno* females had a single UV cell type, females of the *erato/sapho* clade exhibited a novel subdivision into two distinct UV photoreceptor clusters: a female-specific UV1 cluster characterized by high expression of the W-linked *UVRh1* and repression of the autosomal *UVRh2*, and a UV2 cluster characterized by high expression of *UVRh2* (Fig. 6B). Notably, UV2 cells also retained *UVRh1* transcripts; given that previous immunohistochemical studies confirm the proteins are mutually exclusive in the retina^31,32^, we infer that a post-transcriptional mechanism likely blocks the translation of *UVRh1* in UV2 cells to achieve better color discrimination.

We next sought the transcription factors responsible for this bifurcation. Differential expression analysis between female UV1 and UV2 clusters identified *doublesex* (*dsx*), the master regulator of somatic sex differentiation, as the primary candidate (Fig. 6C). We observed a switch-like negative correlation between *dsx* and *UVRh2* expression: high *dsx* levels in UV1 cells correlated with the silencing of the autosomal *UVRh2*, while low *dsx* levels in UV2 cells correlated with the de-repression of *UVRh2*, reverting the cell to a male-like opsin expression profile (Fig. 6D). To determine if this interaction between *UVRh2* and *dsx* was an ancestral feature or a derived innovation, we compared *dsx* and opsin expression patterns in the melpomene/cydno clade. In these species, females ubiquitously expressed high levels of *dsx* in UV photoreceptors, resulting in the constitutive repression of *UVRh2* (Fig. 6E). We further confirmed the absence of UV subtypes in males across all species (Fig. 6B). The emergence of true UV color vision in the *erato/sapho* clade is therefore driven by the redeployment of the sexual differentiation pathway. By evolving a mechanism to stochastically downregulate *dsx* in a subset of photoreceptors, females release the autosomal *UVRh2* from repression, creating a cellular mosaic where some cells maintain female identity while others revert to a male-like transcriptional state.

## Discussion

A central challenge in evolutionary biology is understanding how complex adaptations evolve without disrupting highly conserved functions. With its inherently modular, repeating ommatidia and rapid ecological diversification, the insect compound eye serves as an exceptional system to investigate this evolutionary framework. By integrating population genomics and single-cell transcriptomics of the *Heliconius* compound eye from microevolutionary (intra-population polymorphism) through macroevolutionary (cross-species) timescales, we show that distinct genes and cell types contribute to visual behavior divergence at different stages of evolution.

### Linkage between the color and visual preference genes

In the polymorphic *H. c. alithea*, our *k*-mer GWAS identified *sens-2* as the primary mate preference gene. Evolutionary theory has long proposed that mating cues and their corresponding preferences should be genetically linked under sexual selection^98^. While the mechanisms maintaining this linkage typically involve pleiotropy or tight physical linkage along the linear genome, our findings implicate an additional mechanism: physical proximity in 3D chromatin space. Specifically, Micro-C data revealed long-range contacts between the visual preference gene *sens-2* and the visual cue (wing color) gene *al-1*. This 3D chromatin architecture likely facilitates the co-regulation of these distinct regions. Consequently, mutations could jointly affect both genes, even though *al-1* and *sens-2* are separated by 600 kb on the 2D linear chromosome. Furthermore, we observed numerous other significant genetic variants within the *K* locus, which may also contribute to the complex regulatory landscape governing *sens-2* expression.

### Glia in behavioral evolution

While mate preference genes are often assumed to be neuronal, we unexpectedly found that *sens-2* is uniquely expressed in fenestrated glia, a specialized surface glial cell type at the retina-lamina boundary. Historically relegated to structural and metabolic support roles, glia are now emerging as substrates for behavioral divergence^90,99^. Lamina glia are particularly notable because they are transcriptionally distinct from homologous glial populations found elsewhere in the nervous system^76^. Given their close proximity to photoreceptors, one of their specialized functions is to recycle histamine via a multicellular glial network^100^. Fenestrated glia mediate the final step of this shuttle to the retina, transferring histamine metabolites to pigment cells for subsequent delivery back to the photoreceptors. Furthermore, because *sens-2* expression in surface glia is known to regulate neural development in *Drosophila* larvae^59^, we hypothesize that *sens-2* influences visual mate preference by modulating histamine recycling dynamics or by shaping axon projections during pupal development.

### Evolution of novel photoreceptors

When comparing transcriptomic divergence across multiple species, we found that glia are evolutionarily more conserved than neurons. Among neuronal populations, photoreceptors, particularly the R7 photoreceptors, evolve significantly faster than downstream lamina neurons. This aligns with the hypothesis that peripheral sensory neurons such as photoreceptors are highly labile, providing a flexible substrate for rapid adaptation. Consistent with this accelerated evolution, we identified two novel R7 subtypes: a female-specific UV subtype that originated within the *Heliconius* genus, and a broad-range photoreceptor that likely arose within the Nymphalidae family. Strikingly, both evolutionary innovations arose not through the *de novo* construction of gene regulatory networks, but via the redeployment of existing transcription factors to alter opsin expression. Specifically, the novel UV R7 emerged by downregulating *dsx* in females to accommodate additional male-like UV photoreceptors, whereas the broad-range R7 emerged by downregulating *spalt* to permit long-wavelength opsin expression. Although these represent just two case studies, they illustrate a shared mechanism for generating novel cell types: the relaxation of existing repressor nodes within established regulatory networks to recruit new functions.

Ultimately, the compound eye represents a powerful system for dissecting the multi-scale evolutionary dynamics of cell type evolution. The architecture of the compound eye is fundamentally modular, built from thousands of repeating ommatidia. At short evolutionary time scales, modifying cis-regulatory elements allow for cell-type-specific transcriptome divergence. Over macroevolutionary timescales, novel cell type and novel ommatidia can be incorporated into the retina, and the proportion of these ommatidial subtypes can be further tuned and spatially arranged to fit distinct ecological needs. It is also worth noting that the novel broad-range photoreceptor and associated red color vision can be lost easily. For example, female *Argynnis paphia* and many nymphalid butterfly species are missing the broad-range photoreceptors^101^. The genetic mechanisms leading to the evolution and secondary loss of this novel cell type are still unknown. Future study will need to sample more broadly across the Nymphalidae to fully understand the origin of cell type novelty.

## Data Availability

- Single-nucleus RNA-seq data have been deposited at GEO: GSE330236.
- Single-nucleus ATAC-seq and ATAC-seq data have been deposited at GEO: GSE330347.
- Micro-C data have been deposited at GEO: GSE330237.
- Genome assemblies and re-sequencing data will be deposited at NCBI BioProject PRJNA1460780.
- Any additional information required to reanalyze the data reported in this paper is available upon request.

## Acknowledgement

We thank the University of Chicago Functional Genomics Core Facility (RRID:SCR_019196), Cytometry and Antibody Technology Facility (RRID:SCR_017760), Human Tissue Resource Center (RRID:SCR_004418), and the OBA Light Imaging Facility for research support, as well as the Center for Research Informatics Bioinformatics Core Facility (RRID:SCR_022937) for computational support. We are grateful to Darli Massardo and the University of Chicago greenhouse staff for rearing the butterflies, and to Andres Vega, Luis Ricardo Murillo-Hiller, and Susan D. Finkbeiner for their assistance with fieldwork. We also thank Trevor Price, John Novembre, Stephanie Palmer, Hana Nagata, and Xiao Li for their valuable feedback on the manuscript. Fieldwork was supported in part by a CLACS Tinker Field Research Grant and an OTS Henry Leigh Fellowship awarded to W.L. This work was supported by NIH grant R35 GM118147 to M.S.L. and NIH grant R35 GM131828 to M.R.K.

## Author Contributions

Conceptualization, W.L., N.W.V., and M.R.K.; methodology, W.L., C.C., and N.W.V.; investigation, W.L., N.W.V, C.C., and C.W.; formal analysis, W.L.; data curation, W.L.; writing – original draft, W.L.; writing – review & editing, W.L., N.W.V., C.C., and M.R.K.; supervision, M.R.K.; project administration, M.R.K.; funding acquisition, M.S.L. and M.R.K.

## Methods

### Animals

All commercially obtained butterflies were maintained in greenhouse breeding colonies at the University of Chicago, with stocks periodically supplemented with new individuals. Adult butterflies were provided access to artificial nectar, supplemented with blooming *Lantana* as an additional source of nectar and pollen. Pupae of *Heliconius cydno galanthus* and *H. melpomene* were obtained from El Bosque Nuevo (Costa Rica), and *H. c. alithea* from Heliconius Butterfly Works (Ecuador). Wild specimens, including *H. c. galanthus*, *H. pachinus*, *H. melpomene*, *H. sapho*, and *H. hewitsoni*, were collected and accessed under research permit R-042-2022-OT-CONAGEBIO issued by CONAGEBIO, Costa Rica. All international sample transfers adhered to the legal requirements of the USA, Costa Rica, and Ecuador.

### Nanopore DNA sequencing

High molecular weight (HMW) DNA was extracted from the whole thorax tissue using the QIAGEN Genomic-tip 20/G (#10223). DNA was sequenced on two Oxford Nanopore Technologies (ONT) R9.4.1 flow cells. Library preparation followed a modified version of the *Drosophila* nanopore sequencing protocol^102^. Key modifications included extending the incubation times for both the DNA dA-tailing and adapter ligation steps to 1 hour each. To enrich for long fragments, the final library was size-selected using the Short Read Eliminator (SRE) XS Kit (PacBio, #102-208-200) to remove fragments smaller than 5 kb. Raw signal data were basecalled using Guppy 6.1.5 with the super-accuracy model (dna_r9.4.1_450bps_sup).

### Micro-C experiment

Micro-C libraries were prepared from six adult male *H. c. alithea* brains, including central brains and optic lobes, using a modified low-input protocol optimized for neural tissue^103^. We generated one Micro-C library for white males and one for yellow males. Brains were collected in PBS on ice, fixed in 1% formaldehyde for 15 min at room temperature with gentle mixing, washed twice in PBST for 5 min each, and then subjected to a second crosslinking step in PBST containing 3 mM disuccinimidyl glutarate (DSG) and 3 mM ethylene glycol bis(succinimidyl succinate) (EGS) for 45 min at room temperature with passive mixing. Crosslinking was quenched with 2 M Tris-HCl, pH 7.5, for 5 min, and samples were pelleted by centrifugation for 5 min at 4 °C at maximum speed, washed twice in PBST, snap-frozen in liquid nitrogen, and stored at −80 °C until further processing.

Frozen crosslinked brains were resuspended in 500 µl ice-cold Buffer MB#1 (50 mM NaCl, 10 mM Tris-HCl, pH 7.5, 5 mM MgCl_2_, 1 mM CaCl_2_, 0.2% NP-40, and 1× protease inhibitor cocktail), incubated on ice for 20 min, pelleted briefly, and washed twice in the same buffer. Chromatin was digested in 500 µl Buffer MB#1 with micrococcal nuclease to yield approximately 90% mononucleosomes and 10% dinucleosomes. Digestion was carried out for 10 min at 37 °C with shaking at 900 r.p.m. and stopped by addition of EGTA to a final concentration of 4 mM, followed by incubation for 10 min at 65 °C. Samples were then pelleted and washed twice with cold Buffer MB#2 (50 mM NaCl, 10 mM Tris-HCl, pH 7.5, 10 mM MgCl_2_). Chromatin ends were repaired in situ by sequential treatment with T4 polynucleotide kinase and Klenow fragment in NEBuffer 2.1 supplemented with ATP and DTT for 15 min each at 37 °C. End labeling was then performed in a 75 µl reaction containing biotin-14-dATP, biotin-11-dCTP, dTTP, dGTP, T4 DNA ligase buffer, and BSA for 45 min at 25 °C with intermittent mixing. Reactions were terminated by addition of EDTA to a final concentration of 30 mM and incubation for 20 min at 65 °C, after which samples were pelleted and rinsed three times in cold Buffer MB#3 (50 mM Tris-HCl, pH 7.5, 10 mM MgCl_2_).

For proximity ligation, chromatin pellets were resuspended in a 250 µl ligation mix containing T4 DNA ligase buffer, BSA, and T4 DNA ligase and incubated for 3 hours at 25 °C with shaking at 900 r.p.m. Unligated biotinylated ends were removed by Exonuclease III treatment in NEBuffer 1 for 15 min at 37 °C. Crosslinks were then reversed by addition of SDS and Proteinase K followed by overnight incubation at 65 °C with shaking. DNA was recovered by phenol:chloroform:isoamyl alcohol extraction followed by ethanol precipitation in the presence of sodium acetate and glycogen. Pellets were washed with 75% ethanol, air-dried, and resuspended in EB buffer containing RNase A, followed by incubation for more than 30 min at 37 °C. DNA was further purified using the NucleoSpin Gel and PCR Clean-up kit and eluted in kit elution buffer. Biotinylated ligation products were captured using Dynabeads MyOne Streptavidin C1 beads. Beads were washed in 1× BW buffer, resuspended in 2× BW buffer, and incubated with purified DNA for 20 min at room temperature. After capture, beads were washed twice in 1× BW buffer at 55 °C and once in EB buffer. End repair and A-tailing were then carried out directly on bead-bound DNA using the NEBNext Ultra II DNA End Prep module. Illumina adapters were ligated using the NEBNext Ultra II DNA ligation reagents, followed by USER enzyme treatment. Beads were subsequently washed in 1× BW buffer and EB buffer and resuspended in EB buffer for amplification. Libraries were amplified directly from streptavidin beads using KAPA HiFi HotStart ReadyMix and NEBNext multiplex oligonucleotides with the following cycling conditions: 98 °C for 45 s; 8 cycles of 98 °C for 15 s, 60 °C for 30 s, and 72 °C for 30 s; followed by 72 °C for 1 min. PCR products were size-selected with AMPure XP beads using a two-step procedure in which 0.55× beads were first used to remove fragments larger than 500 bp, followed by an additional 0.4× purification of the supernatant to yield a final 0.95× size selection. Purified libraries were eluted in nuclease-free water and quantified by Qubit fluorometry and Bioanalyzer analysis before sequencing. Libraries were sequenced on a single lane of a NovaSeq X 25B flow cell to generate 150-bp paired-end reads.

### *Heliconius cydno alithea* genome assembly

We assembled the reference genome using ONT reads from a yellow *H. c. alithea* female individual (N29). The GenomeScope^104^ *k*-mer estimated genome size was 282 Mb, identical to the previous estimate of *melpomene*/*cydno*/silvaniform clade genome size^14^. The *k*-mer heterozygosity estimates were high (2.13%-2.24%). We used 282 Mb as the estimated genome size to generate *de novo* assemblies with flye 2.9^105^. We further polished the flye assemblies with two rounds of racon 1.5.0 and one round of medaka 1.7.2 (r941_min_sup_g507_model) and purged haplotigs with purge_dups 1.2.6^106^. To complement the relatively accurate but fragmented flye assemblies, we additionally generated phased diploid assemblies using Shasta 0.13.0^107^ with Nanopore-Phased-May2022 configuration and minimal read length of 10 kb. We set the contiguous shasta haploid assemblies as query sequences and the fragmented purged flye assemblies as reference sequences, merged them with minimum alignment of 10 kb and anchor length cutoff of 500 kb using quickmerge^108^. We did another run of purge_dups after the quickmerge step. Finally, we filtered out contigs smaller than 1 kb since most of these contigs were tandem duplications of the TTAGG telomere motif, which were also found in other Lepidopetra species^109^. Micro-C reads from male *H. c. alithea* brains were mapped to the draft assembly from previous steps using bwa^110^ with SP5M tag. We further filtered valid Micro-C alignments using pairtools^111^ and scaffolded the assembly into a chromosome-scale assembly with YaHS^112^.

Since assemblies derived solely from nanopore reads can have high local error rates, we further polished the assembly with illumina short reads from other *H. c. alithea* individuals and original nanopore long reads (white and yellow females) using NextPolish^113^. The polishing step improved the assembly contiguity. Afterwards, we used CRAQ^114^ to validate the assembly quality and break misassemblies using clipping signals from both short-read and long-read alignments. We also filtered contigs without 99% coverage. Finally, we renamed and oriented each chromosome according to the *Heliconius melpomene*^115^ (Hmel2.5) reference genome. The mitochondrial genome was assembled with GetOrganelle^116^ using illumina short reads from a different *H. c. alithea* individual.

### TAD, loop, and compartment identification from Micro-C

Micro-C paired-end sequencing reads were aligned to the reference genome using BWA-MEM2 2.2.1 with the SP5M flag. The resulting alignments were parsed, sorted, and deduplicated using pairtools 1.1.2. During the initial parsing step, alignments mapping to the mitochondrial genome were filtered out. Pairs were further filtered to retain only uniquely mapped and uniquely rescued contacts (UU, UR, and RU pair types). The final valid interaction pairs were extracted using pairtools split and indexed utilizing pairix. Contact matrices were subsequently constructed from these valid pairs at a 100-base-pair resolution using cooler 0.10.3 cload.

TADs were identified using hicFindTADs from HiCExplorer 3.7.5 on Knight-Ruiz (KR) normalized contact matrices at both 5 kb and 15 kb resolutions. minDepth was set at 3x and maxDepth at 10x resolution size. Loops were identified using Mustache 1.0.1 on KR-normalized contact matrices at 1.5 kb with the p-value threshold of 0.01. Compartments were identified using PC2 from hicPCA (HiCExplorer 3.7.5). A (active) and B (inactive) compartments were further assigned based on the gene expression level from bulk RNA-seq.

### ATAC-seq

We performed ATAC experiments following published protocol^61,117^ with minor modifications. Brains were dissected in room temperature PBS then immediately transferred into ice cold sucrose buffer with protease inhibitor (10 mM Tris-HCl, 250 mM D-sucrose, 1 mM MgCl_2_, pH 7.5) in 1 mL dounce homogenizers. Tissues were dissociated using 60 strokes with the tight pestle, then transferred to cold 1.5 mL tubes. Cells and nuclei were pelleted by centrifugation for 5 min at 1000 x g at 4 °C, then resuspended in 150 uL lysis buffer (wash buffer: 10 mM Tris-HCl pH 7.5, 10 mM NaCl, 3 mM MgCl_2_ plus 0.2% NP-40) and lysed on ice for 5 min. Nuclei were harvested by centrifugation for 5 min at 1000 x g for 5 min and 4 °C, then resuspended in 750 uL ice cold wash buffer and stained with Hoechst for cell counting. Aliquots of 500,000 cells were pelleted for 7 min at 1000 x g and 4 °C, then resuspended in transposition mix (25 uL TD buffer, 2.5 uL TDE1 enzyme, 22.5 uL water). Transposition was performed 30 min at 37 °C shaking at 1000 rpm, then cleaned up using the Zymo DNA Clean and Concentrator 5 kit. Libraries were amplified 10-11 cycles before double-sided cleanup (0.5X – 1.8X) with SPRI Select Beads (Beckman-Coulter, USA). We sequenced libraries as 50 bp paired-end reads on a single lane of a NovaSeq X 10B flowcell, yielding an average of 30M read pairs per sample. We followed the ENCODE ATAC-seq pipeline^118^ to process raw sequencing reads.

### Nanopore cDNA-PCR sequencing

To improve our genome annotation, we performed Oxford Nanopore cDNA-PCR sequencing on *H. c. alithea* adult retina tissue, as well as pupal (96h post-pupation) brain, retina, and forewing tissues (Table S8). RNA was isolated via phenol-chloroform extraction, and the RNA pellet was precipitated using glycogen and –20 °C isopropanol. Following extraction, cDNA libraries were prepared using the cDNA-PCR Sequencing Kit (SQK-PCS111, ONT) and the PCR-cDNA Barcoding Kit (SQK-PCB111.24, ONT) according to the manufacturer’s protocol. The final libraries were sequenced across three R9.4.1 flow cells. Raw sequencing signals were basecalled using the dna_r9.4.1_e8_sup@v3.6 model in Dorado. We filtered the output to retain reads with a mean Phred quality score ≥ 7, yielding a total of 37 million reads with an N50 of 1.2 kb. To orient the reads and trim adapters, we used Pychopper 2.7.10 to identify and rescue full-length cDNA reads. Approximately 70% of the sequences were full-length cDNA with strand-switching primers and VN primers in the correct orientation, which we designated as stranded transcripts. The remaining reads were classified as unstranded.

### Long-read stringtie assembly

ONT cDNA reads from various tissues were pooled and mapped to the *H. c. alithea* reference genome using the spliced long-read preset (-ax spliced) in minimap2 2.29^119^. Stranded reads identified via Pychopper and unstranded reads were aligned in separate runs. The resulting BAM files were subsequently used to construct independent stranded and unstranded long-read assemblies using StringTie with the long-read option (-L).

### Short-read stringtie assembly

We downloaded previously published developing brain *H. c. alithea* bulk RNA-seq datasets (PRJNA1019262)^34^. Reads were aligned to the reference genome using HISAT 2.2.1^120^ with the –-dta flag optimized for downstream transcriptome assembly. Following alignment, we generated strand-specific short-read assemblies using StringTie 3.0.0^121^. To ensure high-confidence transcript models, we applied strict coverage thresholds, requiring a minimum of 3x coverage for multi-exon transcripts, 5x coverage for single-exon transcripts, and a minimum assembled transcript length of 200 bp. To minimize the assembly of noisy isoforms and artifacts from low-abundance transcription, we required a minimum isoform fraction of 20%, at least five supporting reads per splice junction, and a minimum junction anchor length of 15 bp. Additionally, we capped the allowable multi-mapping fraction per locus at 50%. Finally, to account for the compact gene arrangement, we reduced the maximum allowed gap between read mapping to 25 bp. To specifically address potential read-through transcripts, we generated an alternative consensus transcriptome using TACO 0.7.3^122^, which leverages expression abundance to accurately demarcate transcript boundaries.

### Prediction of protein coding genes with BRAKER3

To identify coding sequences (CDS) and rescue lowly expressed genes potentially missed by transcriptome assembly, we performed protein-coding gene prediction using the BRAKER3^123^ pipeline. We supplied *H. c. alithea* brain RNA-seq data and the Arthropoda partition of OrthoDB v12 dataset^124^, which was aligned using DIAMOND^125^ and Spaln^126^. This combined transcriptomic and protein evidence was used to iteratively train GeneMark-ETP^127^ and AUGUSTUS^128^. Finally, TSEBRA^129^ evaluated the candidate models against the extrinsic evidence to output the optimal annotations.

### Transcript selection using Mikado

We generated a final transcriptome by integrating multiple independent assemblies using Mikado 2.3.4^130^. The input GTF assemblies comprised annotations transferred via Liftoff 1.6.3^131^ from both the *H. c. chioneus* assembly^132^ and a prior *H. c. alithea* assembly^34^, alongside StringTie short-read assemblies, long-read assemblies, and BRAKER3 gene predictions. TransDecoder was used to identify open reading frames (ORFs) within the StringTie assemblies, and transcripts were aligned against the Arthropoda OrthoDB v12 database using DIAMOND BLASTx. Permissive chimera-splitting parameters were applied to filter readthrough artifacts based on the BLAST results. To accommodate genes with massive introns, the maximum intron size was set to 350 kb. Transcript selection incorporated additional external evidence such as expression abundance (TPM) from Salmon 1.10.1^133^ and coding potential from CPC2 1.0.1^134^. Crucially, the Mikado scoring configuration was adjusted to penalize transcript models lacking complete untranslated regions (UTRs), yielding high-fidelity annotations optimized for downstream snRNA-seq analysis. Mitochondrial genes were annotated separately with MITOS2^135^.

### Ortholog inference and functional annotation

To accurately identify homologs with *Drosophila melanogaster* proteins, we ran OrthoFinder v3.1.0^136^ using our new *H. c. alithea* and *H. sapho* annotations alongside *Drosophila melanogaster* (BDGP6.54), and several Lepidopteran and Dipteran reference proteomes downloaded from Ensembl Metazoa and Ensembl Rapid Release to break up tree branches for accurate inference of orthologs. These additional reference sets included primary protein transcripts from *Anopheles gambiae* (AgamP4), *Bombyx mori* (Bmori_2016v1.0), *Danaus plexippus* (Dplex_v4), *Heliconius melpomene* (Hmel1), *Pieris napi* (GCA_905475465.2), *Spodoptera frugiperda* (AGI_APGP_CSIRO_Sfru_2.0), and *Vanessa atalanta*

(GCA_905147765.2). To standardize nomenclature, *Heliconius* genes with one-to-one orthologs in *Drosophila melanogaster* were renamed to match their respective FlyBase symbols. In cases of non-unique mappings (e.g., one-to-many or many-to-one), *Heliconius* genes were annotated using one of the mapped *Drosophila* symbols followed by the *Heliconius* chromosome and unique gene identifier to ensure unambiguous naming. We further annotated the protein-coding genes using eggNOG-mapper 2.1.13 and the eggNOG database 5.0.2^137^, incorporating Pfam realignment.

### SNP-based GWAS

Whole-genome re-sequencing data for *H. c. alithea* were obtained from BioProject PRJNA802836. While our general pipeline for the SNP-based GWAS followed the methods in our previous work^34^, we introduced several modifications to the data processing steps. Most notably, sequencing reads were mapped to the newly assembled *H. c. alithea* reference genome. Additionally, our quality control parameters were updated: we retained 109 of the 113 sequenced individuals, set genotype calls with a genotype quality (GQ) < 5 and depth (DP) < 1 as missing, and excluded sites with a missing data rate exceeding 10% across individuals. Unlike our previous study, we did not perform linkage-disequilibrium (LD) pruning. After applying these filters, a total of 7 million SNPs were retained for downstream analysis.

We maintained the mate-choice modeling framework from our previous study^34,41^, analyzing courtship events via a generalized linear mixed model (GLMM) implemented in GMMAT^46^. The response variable (whether a male chose a white female or not) was modeled as a binary outcome using a logit link function, treating distinct courtship events as repeated measurements for each individual male. To appropriately control for confounding structure, the kinship matrix, male ID, and specific mating trial were included as random effects. The presence of a yellow female with a triangle melanin patch was included as a fixed effect, as previous research^34^ demonstrates males generally avoid this phenotype combination. Finally, we performed Wald tests across all 7 million retained SNPs.

### *k-mer* GWAS

To construct a binary *k-mer* presence/absence genotype matrix, we utilized the kGWASflow^138^ Snakemake pipeline, which was based on the kmersGWAS^45^ library. Canonical and non-canonical 25-mers were counted from adapter-trimmed reads using KMC^139^. We then applied a strict filtering threshold, retaining only *k-mers* present in more than 5% of the individuals. These were merged into a PLINK-formatted presence/absence genotype matrix, yielding a final set of 126 million unique *k-mers* for downstream analysis.

The *k-mer*-based GWAS was conducted using GMMAT similar to our SNP-based analysis, and the previously calculated SNP-based kinship matrix was used to control for population structure. To optimize computational efficiency across the 126 million *k-mers*, we implemented a two-tiered testing strategy in GMMAT. First, a computationally lightweight score test was performed on all *k-mers*. 193,860 *k-mers* that passed a preliminary significance threshold (p < 0.001) were subsequently subjected to a more rigorous, computationally intensive Wald test. A final significance threshold was determined using the FDR 0.01 threshold established in the SNP-based GWAS (p < 1.86 × 10^-8^).

Because some *k-mers* originate from structural variants (such as insertions) absent from the reference genome, we assembled *k-mer*-containing reads into longer contigs prior to reference mapping. For each individual, reads containing significant *k-mers* were extracted using seal from BBTools 39.0.1^140^. These reads were then pooled across all samples and *de novo* assembled using tadpole from BBTools, producing extended *k-mer* sequences ranging from 124 to 912 bp in length.

These extended contigs were aligned to the reference genome using Bowtie2. Recognizing that a single *k-mer* could be contained in multiple extended sequences, we filtered *k-mer* alignment results to eliminate *k-mer* originating from transposable elements or repetitive regions. First, any *k-mer* whose extended sequences mapped to multiple chromosomes was discarded. Second, *k-mers* were excluded if the maximum genomic distance between their associated contigs exceeded 1,000 bp. For the uniquely anchored *k-mers*, a single representative genomic coordinate was designated using the minimum start position among the contigs, and mapping confidence was reported as the average mapping quality score across all hits.

### Adult eye dissociation

Because the butterfly compound eye features a fused rhabdom and is encapsulated in an exoskeleton cuticle, it is largely inaccessible for standard single-cell RNA sequencing. We therefore performed single-nucleus RNA sequencing (snRNA-seq) by adapting established *Drosophila* snRNA-seq protocols^141^. Adult butterfly eyes were dissected under a stereomicroscope in ice-cold 1x PBS. Dissected eyes were placed in 1.5 ml Eppendorf tubes fully immersed in 200 ul of 1x PBS and immediately snap-frozen in liquid nitrogen. Field-collected samples were maintained in a liquid nitrogen dry shipper following snap-freezing and subsequently stored at –80 °C at the University of Chicago until library preparation.

On the day of library preparation, snap-frozen samples were thawed on ice. Once thawed, 3-6 eyes were transferred to a chilled 1 ml Dounce homogenizer containing 1 ml of freshly prepared homogenization buffer (250 mM Sucrose, 10 mM Tris pH 8.0, 25 mM KCl, 5 mM MgCl2, 0.1% Triton X-100, 0.5% RNase inhibitor [NEB #M0628], 1x protease inhibitor, and 0.1 mM DTT). Nuclei were isolated by douncing 20 times with a loose-fitting pestle followed by 40 times with a tight-fitting pestle. The resulting homogenate was filtered through a 40 um cell strainer (Bel-Art Flowmi) and centrifuged at 1000 x g for 10 minutes at 4°C. The nuclei pellet was resuspended in 500 ul of resuspension buffer (1x PBS, 0.5% BSA, 0.5% RNase inhibitor) and filtered a second time using a 20 um pluriStrainer.

To eliminate cell clumps and debris, nuclei were stained with Hoechst and diluted with an additional 500 ul of resuspension buffer. Nuclei were sorted using a BD FACSAria III Fusion flow cytometer equipped with a 100 um nozzle. Hoechst-positive singlets were collected into resuspension buffer, with clumps and doublets excluded via standard gating strategies. Following sorting, the nuclei underwent a final wash (centrifugation at 1000 x g for 10 minutes at 4 °C) and the pellet were resuspended in 100 ul of resuspension buffer. This final suspension was submitted to the University of Chicago Genomics Facility for library preparation using Chromium Next GEM Single Cell 3’ Reagent Kit v3.1 kit (10x Genomics).

### Libraries were sequenced with NovaSeq X. snRNA-seq quality control and preprocessing

Sequencing FASTQ files were aligned to custom *Heliconius* reference genomes and annotations using Cell Ranger 10.0.0 with intronic reads included (Table S3). Reads from *H. melpomene*, *H. cydno*, and *H. pachinus* were mapped to the *H. cydno alithea* reference genome, while reads from *H. sapho* and *H. hewitsoni* were mapped to the *H. sapho* reference.

To distinguish cell-containing droplets from empty droplets and remove ambient RNA contamination, the raw gene count matrices produced by Cell Ranger were processed using CellBender^142^ 0.3.0. The software was run on a GPU node for 150 epochs with a learning rate of 0.000025 and a false positive rate of 0.05.

To demultiplex pooled individuals and identify doublets, we utilized the Demuxafy 3.0.0 pipeline. Demuxalot^143^ 0.4.3 with refinement was first used to assign nuclei to specific individuals based on reference genotypes and to identify heterotypic genotype doublets, which are formed by cells from different individuals. We then performed transcriptome-based doublet detection using scDblFinder^144^ 1.18.0, incorporating the doublets identified by Demuxalot as known priors to improve sensitivity.

To estimate the global doublet rate, including both homotypic and heterotypic doublets, we employed a custom R script (fit_doublets.R). By assuming a Poisson distribution for the number of nuclei in a droplet, the model optimized the joint likelihood of observed singlets and heterotypic doublets to infer the frequency of hidden homotypic doublets. Parameter optimization was achieved using the BFGS algorithm. Finally, we excluded all droplets identified as doublets by either Demuxalot or scDblFinder, as well as those with a mitochondrial gene proportion greater than 1%.

### Cross library snRNA-seq integration

snRNA-seq data processing was done in Seurat 5.4.0. To select for highly variable genes (HVGs), we first used ROGUE^65^ to identify 979 variable genes with a significance threshold of p.adjust < 0.1 in the *H. c. alithea* dataset. Following log-normalization of the expression data, the first 30 principal components (PCs) were calculated based on these selected HVGs. To account for batch effects, the filtered snRNA-seq data were then integrated by individual assignment using Harmony^68^. Finally, unsupervised cell clustering was performed on the integrated data using the Leiden algorithm at a resolution of 0.5.

The final integrated atlas was subjected to cluster-level filtering. First, to remove clusters likely consisting of ambient RNA droplets, we excluded any cluster with a median of fewer than 300 detected genes (median nFeature_RNA < 300). Second, clusters representing potential dissection artifacts were removed if they were heavily biased by a single biological replicate, defined as any cluster in which over 50% of the cells originated from one individual.

### scATAC-seq data

To generate a single scATAC-seq library, we pooled six eyes from 3-day-old adult male *H. c. galanthus*. Nuclei suspensions were prepared following the snRNA-seq protocol described above. The final suspension was submitted to the University of Chicago Genomics Facility for library preparation using the Chromium Next GEM Single Cell ATAC v1.1 kit (10x Genomics). The library was sequenced on a single lane of the NovaSeq X 10B flow cell. Raw sequencing reads were mapped to the *H. c. alithea* reference genome and annotated using Cell Ranger ATAC v2.2.0.

Cell Ranger outputs were further processed using Signac^81^. We initially recovered 7,570 nuclei (median 4,540 high-quality fragments per cell) and retained those meeting the following quality thresholds: 1,000– 30,000 peak region fragments, >15% of reads in peaks, nucleosome signal <4, and transcription start site (TSS) enrichment >1.5. Following quality control, dimensionality reduction was performed via Latent Semantic Indexing (LSI; utilizing TF-IDF normalization followed by SVD), and UMAP was computed using LSI dimensions 2–30. To assign cell-type identities, we calculated a log-normalized imputed gene activity matrix to identify Canonical Correlation Analysis (CCA) integration anchors, allowing us to transfer labels directly from our pre-processed *H. c. alithea* snRNA-seq reference.

Finally, to resolve cell-type-specific chromatin accessibility, nuclei were grouped by their newly predicted cell type labels for *de novo* peak calling with MACS2. The original fragment data were then re-quantified against this refined MACS2 peak set to construct the final set of chromatin peaks for downstream analyses. To identify differentially accessible regions across cell types, we performed differential accessibility (DA) analysis on the MACS2 peak assay using the FindAllMarkers function from Seurat. DA peaks were retained if they exhibited a log-fold change greater than 1 and were detected in at least 5% of cells.

### Gene expression evolution analysis

We first used AggregateExpression in Seurato 5.4.0 to generate pseudobulk gene count matrices for each individual and cell type combination. We then calculated counts per million (CPM) in the aggregated pseudobulk data to normalize the count. We modeled gene expression evolution using CAGEE with the inferred population tree (Supplementary Fig. 11). Each population had 3-9 replicates. We fitted multiple rates across cell type groups. To control for sampling variation within each branch, we restricted our analysis to common cell types present in at least three individuals across all six populations. Any individuals with less than 5 cells for any of the common cell types were filtered out.

### *sens-2* HCR

We designed HCR probe sets for target genes using HCRProbeMakerCL^145^. Specificity of probe pairs was checked using BLASTn. Probe pairs with matches to non-target transcripts (E-value < 1e-10) were excluded from final probe sets. Probe sets were ordered as 50 pmol oPools from IDT (USA) and resuspended to 1 uM in 10 mM Tris-HCl pH 8.0 before use.

Adult sections were prepared from fresh/frozen tissues. Snap frozen eyes and brains were sectioned at 10 um in a cryostat and stored on slides at –80 °C until use. HCR was performed following Choi et al. (2018) using buffers and reagents purchased from Molecular Instruments (USA). Frozen sections were incubated at 37 °C for 1 min to affix to slides, then fixed for 15 min in ice-cold 4% formaldehyde in PBS. Slides were then dehydrated 5 min each in 50% ethanol in PBS, 70% ethanol in PBS, 100% ethanol, and 100% ethanol at room temperature (RT). Sections were digested 10 min at RT with 10 ug/mL proteinase K in PBS, then washed 2 x 5 min in PBS. Sections were pre-hybridized in Probe Hybridization Buffer for 30 min at 37 °C. Hybridization solutions comprised 4 nM each probe set. Hybridization was carried out overnight at 37 °C. Slides were washed (15 min each at 37 °C) with Probe Wash Buffer (PWB), 75% PWB/25% 5X SSCT, 50% PWB/5X SSCT, 25% PWB/ 5X SSCT, and 5X SSCT. After a final 5 min wash in RT 5X SSCT, slides were pre-amplified in Amplification Buffer for 30 min at RT. Hairpins were heated and snap-cooled separately. Hairpins were used at 6 nM. Amplification proceeded overnight. Slides were washed for 5 min in 5X SSCT at RT, 30 min in 5X SSCT + 1 ug/mL DAPI, 30 min in 5X SSCT, and a final 5 min in 5X SSCT before being mounted in Vectashield. Slides were imaged on a Zeiss LSM 900 or LSM 710 in the University of Chicago OBA Light Imaging Facility.

### *sens-2* CRISPR/Cas9 knockouts

We attempted to knock out *sens-2* by using CRISPR/Cas9 to induce small frameshifts near the start codon. We identified CRISPR target sites within *sens-2* exon 1 using IDT’s (USA) crRNA Design Tool, then synthesized sgRNAs and performed injections following^36^. Initial high-resolution melt analyses on eggs two days after injection suggested high mutation efficiencies (30% – 100%), but hatching rates were consistently low (<10%) and survival to pupation even lower (<5%). These rates were consistent across a number of gRNAs, combinations of gRNAs, and concentrations of RNP (Table S6-S7).

### *Drosophila melanogaster sens-2* staining

We raised a new polyclonal antibody in rabbits against the *Drosophila melanogaster* sens-2 peptide antigen RRDEPPPQRRTPRS at GenScript (NJ, USA). We then used this antibody to stain for sens-2 in the adult *Drosophila* retina following standard protocols. Preliminary stains were performed in adult Oregon-R brains and retinas dissected from flies less than three days old. Some stains included anti-Pros antibody (MR1A) available from the Developmental Hybridoma Studies Databank (USA). Antibodies were used at 1:100 dilutions.

### *Heliconius* genome re-sequencing and variant calling

Genomic DNA was isolated from the thorax of *Heliconius* individuals used in our snRNA-seq analysis using chloroform extractions. Illumina paired-end libraries were constructed using the KAPA Hyper Prep Kit (KAPA Biosystems) or Nextera Library Prep Kit and sequenced to ∼30X using 150 bp paired-end reads on an Illumina HiSeq 2500 or HiSeq 4000 at the University of Chicago Genomics Facility. Low-quality regions and adapters were trimmed from raw reads using Trimmomatic before mapping to the *H. c. alithea* reference using bowtie2 v2.3.2 with default settings except –-very-sensitive-local. We then marked PCR duplicate reads with Picard and realigned around putative indels using the Genome Analysis Toolkit (GATK) 3.8. SNP and indel calling were performed using the GATK’s UnifiedGenotyper module with the heterozygosity priors set to 0.02 and 0.0002 for SNPs and indels, respectively.

### Phylogenetic tree construction

SNPs were rigorously filtered using BCFtools. We retained biallelic SNPs with a minimum site quality (QUAL) of 30. Individual genotypes lacking sufficient support, defined as a read depth (DP) < 10 or a genotype quality (GQ) < 30, were converted to missing data. After recalculating allele frequencies, we required each retained site to have at least one valid allele call (AN > 0) and a maximum missing data frequency of 20%. The filtered VCF was converted to PHYLIP format for phylogenetic inference. We used CASTER-SITE^146^, incorporating branch lengths, to infer relationships among individuals and reconstruct the overall population phylogeny. The result for population phylogeny is shown below. The population tree branch length is in million years.

Population phylogeny newick: ((pacific_melpomene:1.351912004,caribbean_melpomene:1.351912004):0.7006334851,(alithea:1.57128 8465,(captive_bred_galanthus:1.460673557,(pachinus:1.332427859,galanthus:1.332427859):0.1282456 986):0.110614908):0.4812570237);

### *Heliconius sapho* genome assembly and annotation

For our analyses, we used the *Heliconius sapho* reference genome assembly ilHelSaph4 (INSDC Assembly Accession: GCA_964147245.1; BioProject: PRJEB76212), generated by the Wellcome Sanger Institute Tree of Life Programme^147^. To account for the missing W chromosome in this male-derived assembly, we appended the *H. hewitsoni* W chromosome^21^ to the *H. sapho* genome prior to downstream analysis. The mitochondrial genome was independently assembled using GetOrganelle with Illumina short reads from an *H. sapho* individual.

Transcriptome assembly and annotation followed a pipeline similar to our *H. c. alithea* approach. We obtained *H. sapho* RNA-seq data from BioProject PRJEB68281, and used Mikado to select the optimal consensus transcripts from a combination of StringTie short-read transcript assemblies, BRAKER3 gene predictions, and transferred annotations from *H. hewitsoni*^21^.

## Notes

### Competing Interest Statement

The authors have declared no competing interest.

### Summary of Updates

Author affiliations and ORCID updated

## References

1. Nilsson, D.-E. (2013). Eye evolution and its functional basis. Visual Neurosci. 30, 5–20. 10.1017/S0952523813000035.

2. Bowmaker, J.K. (2008). Evolution of vertebrate visual pigments. Vision Research 48, 2022–2041. 10.1016/j.visres.2008.03.025.

3. McCulloch, K.J., Macias-Muñoz, A., and Briscoe, A.D. (2022). Insect opsins and evo-devo: what have we learned in 25 years? Philosophical Transactions of the Royal Society B: Biological Sciences 377. 10.1098/rstb.2021.0288.

4. Briscoe, A.D., and Chittka, L. (2001). The evolution of color vision in insects. Annual Review of Entomology 46, 471–510. 10.1146/annurev.ento.46.1.471.

5. van der Kooi, C.J., Stavenga, D.G., Arikawa, K., Belušič, G., and Kelber, A. (2021). Evolution of Insect Color Vision: From Spectral Sensitivity to Visual Ecology. Annual Review of Entomology 66, 435–461. 10.1146/annurev-ento-061720-071644.

6. Hahn, J., Monavarfeshani, A., Qiao, M., Kao, A.H., Kölsch, Y., Kumar, A., Kunze, V.P., Rasys, A.M., Richardson, R., Wekselblatt, J.B., et al. (2023). Evolution of neuronal cell classes and types in the vertebrate retina. Nature 624, 415–424. 10.1038/s41586-023-06638-9.

7. Wang, J., Zhang, L., Cavallini, M., Pahlevan, A., Sun, J., Morshedian, A., Fain, G.L., Sampath, A.P., and Peng, Y.-R. (2024). Molecular characterization of the sea lamprey retina illuminates the evolutionary origin of retinal cell types. Nat. Commun. 15, 10761. 10.1038/s41467-024-55019-x.

8. Gavriouchkina, D., Tan, Y., Parey, E., Ziadi-Künzli, F., Hasegawa, Y., Piovani, L., Zhang, L., Sugimoto, C., Luscombe, N., Marlétaz, F., et al. (2025). A single-cell atlas of the bobtail squid visual and nervous system highlights molecular principles of convergent evolution. Nat. Ecol. Evol. 9, 1245– 1262. 10.1038/s41559-025-02720-9.

9. Duruz, J., Sprecher, M., Kaldun, J.C., Al-Soudy, A.-S., Lischer, H.E., van Geest, G., Nicholson, P., Bruggmann, R., and Sprecher, S.G. (2023). Molecular characterization of cell types in the squid loligo vulgaris. eLife 12, e80670. 10.7554/eLife.80670.

10. Yeung, K., Bollepogu Raja, K.K., Shim, Y.K., Li, Y., Chen, R., and Mardon, G. (2022). Single cell RNA sequencing of the adult drosophila eye reveals distinct clusters and novel marker genes for all major cell types. Commun. Biol. 2022 5:1 5, 1–18. 10.1038/s42003-022-04337-1.

11. Özel, M.N., Simon, F., Jafari, S., Holguera, I., Chen, Y.C., Benhra, N., El-Danaf, R.N., Kapuralin, K., Malin, J.A., Konstantinides, N., et al. (2021). Neuronal diversity and convergence in a visual system developmental atlas. Nature 589, 88–126. 10.1038/S41586-020-2879-3.

12. Li, H., Janssens, J., De Waegeneer, M., Kolluru, S.S., Davie, K., Gardeux, V., Saelens, W., David, F.P.A., Brbić, M., Spanier, K., et al. (2022). Fly cell atlas: a single-nucleus transcriptomic atlas of the adult fruit fly. Science 375, eabk2432. 10.1126/science.abk2432.

13. Kozak, K.M., Wahlberg, N., Neild, A.F.E., Dasmahapatra, K.K., Mallet, J., and Jiggins, C.D. (2015). Multilocus species trees show the recent adaptive radiation of the mimetic heliconius butterflies. Syst. Biol. 64, 505–524. 10.1093/sysbio/syv007.

14. Cicconardi, F., Milanetti, E., Pinheiro de Castro, E.C., Mazo-Vargas, A., Van Belleghem, S.M., Ruggieri, A.A., Rastas, P., Hanly, J., Evans, E., Jiggins, C.D., et al. (2023). Evolutionary dynamics of genome size and content during the adaptive radiation of heliconiini butterflies. Nat. Commun. 14, 5620. 10.1038/s41467-023-41412-5.

15. Jiggins, C.D., Naisbit, R.E., Coe, R.L., and Mallet, J. (2001). Reproductive isolation caused by colour pattern mimicry. Nature 411, 302–305. 10.1038/35077075.

16. Kronforst, M.R., Young, L.G., Kapan, D.D., McNeely, C., O’Neill, R.J., and Gilbert, L.E. (2006). Linkage of butterfly mate preference and wing color preference cue at the genomic location of wingless. Proc. Natl. Acad. Sci. U. S. A. 103, 6575–6580. 10.1073/pnas.0509685103.

17. Merrill, R.M., Gompert, Z., Dembeck, L.M., Kronforst, M.R., McMillan, W.O., and Jiggins, C.D. (2011). Mate preference across the speciation continuum in a clade of mimetic butterflies. Evolution 65, 1489–1500. 10.1111/j.1558-5646.2010.01216.x.

18. Kronforst, M.R., Young, L.G., and Gilbert, L.E. (2007). Reinforcement of mate preference among hybridizing heliconius butterflies. J. Evol. Biol. 20, 278–285. 10.1111/j.1420-9101.2006.01198.x.

19. Rosser, N., Seixas, F., Queste, L.M., Cama, B., Mori-Pezo, R., Kryvokhyzha, D., Nelson, M., Waite-Hudson, R., Goringe, M., Costa, M., et al. (2024). Hybrid speciation driven by multilocus introgression of ecological traits. Nature 628, 811–817. 10.1038/s41586-024-07263-w.

20. Finkbeiner, S.D., and Briscoe, A.D. (2021). True UV color vision in a female butterfly with two UV opsins. J. Exp. Biol. 224. 10.1242/jeb.242802.

21. Chakraborty, M., Lara, A.G., Dang, A., McCulloch, K.J., Rainbow, D., Carter, D., Ngo, L.T., Solares, E., Said, I., Corbett-Detig, R.B., et al. (2023). Sex-linked gene traffic underlies the acquisition of sexually dimorphic UV color vision in heliconius butterflies. Proc. Natl. Acad. Sci. 120, e2301411120. 10.1073/pnas.2301411120.

22. Moura, P.A., Young, F.J., Monllor, M., Cardoso, M.Z., and Montgomery, S.H. (2023). Long-term spatial memory across large spatial scales in heliconius butterflies. Curr. Biol. 33, R797–R798. 10.1016/j.cub.2023.06.009.

23. Dell’Aglio, D.D., Losada, M.E., and Jiggins, C.D. (2016). Butterfly learning and the diversification of plant leaf shape. Front. Ecol. Evol. 4, 1–7. 10.3389/fevo.2016.00081.

24. Williams, K.S., and Gilbert, L.E. (1981). Insects as selective agents on plant vegetative morphology: egg mimicry reduces egg laying by butterflies. Science 212, 467–469. 10.1126/science.212.4493.467.

25. Finkbeiner, S.D., Briscoe, A.D., and Reed, R.D. (2014). Warning signals are seductive: relative contributions of color and pattern to predator avoidance and mate attraction in heliconius butterflies. Evolution 68, 3410–3420. 10.1111/evo.12524.

26. Couto, A., Young, F.J., Atzeni, D., Marty, S., Melo-Flórez, L., Hebberecht, L., Monllor, M., Neal, C., Cicconardi, F., McMillan, W.O., et al. (2023). Rapid expansion and visual specialisation of learning and memory centres in the brains of heliconiini butterflies. Nat. Commun. 14, 4024. 10.1038/s41467-023-39618-8.

27. Wright, D.S., Rodriguez-Fuentes, J., Ammer, L., Darragh, K., Kuo, C.-Y., McMillan, W.O., Jiggins, C.D., Montgomery, S.H., and Merrill, R.M. (2024). Selection drives divergence of eye morphology in sympatric heliconius butterflies. Evolution 78, 1338–1346. 10.1093/evolut/qpae073.

28. Seymoure, B.M., Seymoure, B.M., Mcmillan, W.O., and Rutowski, R. (2015). Peripheral eye dimensions in longwing (heliconius) butterflies vary with body size and sex but not light environment nor mimicry ring. J. Res. Lepid. 48, 83--92. 10.5962/p.266475.

29. Dang, A., Cerkvenik, U., Ilić, M., Pirih, P., Debevc, E., Briscoe, A.D., and Belušič, G. (2025). Graded opsin co-expression along the butterfly retina fine tunes the spectral sensitivity of a colour-opponent cell across the visual field. J. Comp. Physiol. a, 1–14. 10.1007/s00359-025-01761-6.

30. Briscoe, A.D., Bybee, S.M., Bernard, G.D., Yuan, F., Sison-Mangus, M.P., Reed, R.D., Warren, A.D., Llorente-Bousquets, J., and Chiao, C.C. (2010). Positive selection of a duplicated UV-sensitive visual pigment coincides with wing pigment evolution in heliconius butterflies. Proc. Natl. Acad. Sci. U. S. A. 107, 3628–3633. 10.1073/pnas.0910085107.

31. McCulloch, K.J., Yuan, F., Zhen, Y., Aardema, M.L., Smith, G., Llorente-Bousquets, J., Andolfatto, P., and Briscoe, A.D. (2017). Sexual dimorphism and retinal mosaic diversification following the evolution of a violet receptor in butterflies. Mol. Biol. Evol. 34, 2271–2284. 10.1093/MOLBEV/MSX163.

32. McCulloch, K.J., Osorio, D., and Briscoe, A.D. (2016). Sexual dimorphism in the compound eye of heliconius erato: a nymphalid butterfly with at least five spectral classes of photoreceptor. J. Exp. Biol. 219, 2377–2387. 10.1242/jeb.136523.

33. McCulloch, K.J., Macias-Muñoz, A., Mortazavi, A., and Briscoe, A.D. (2022). Multiple mechanisms of photoreceptor spectral tuning in heliconius butterflies. Mol. Biol. Evol. 39. 10.1093/molbev/msac067.

34. VanKuren, N.W., Buerkle, N.P., Lu, W., Westerman, E.L., Im, A.K., Massardo, D., Southcott, L., Palmer, S.E., and Kronforst, M.R. (2025). Genetic, developmental, and neural changes underlying the evolution of butterfly mate preference. PLOS Biol. 23, e3002989. 10.1371/journal.pbio.3002989.

35. Buerkle, N.P., VanKuren, N.W., Westerman, E.L., Kronforst, M.R., and Palmer, S.E. (2025). Sex-limited diversification of the eye in Heliconius cydno butterflies. J Comp Physiol A. 10.1007/s00359-025-01768-z.

36. Perry, M., Kinoshita, M., Saldi, G., Huo, L., Arikawa, K., and Desplan, C. (2016). Molecular logic behind the three-way stochastic choices that expand butterfly colour vision. Nature 535, 280–284. 10.1038/nature18616.

37. Wernet, M.F., Mazzoni, E.O., Çelik, A., Duncan, D.M., Duncan, I., and Desplan, C. (2006). Stochastic spineless expression creates the retinal mosaic for colour vision. Nature 440, 174–180. 10.1038/nature04615.

38. Anderson, C., Reiss, I., Zhou, C., Cho, A., Siddiqi, H., Mormann, B., Avelis, C.M., Deford, P., Bergland, A., Roberts, E., et al. (2017). Natural variation in stochastic photoreceptor specification and color preference in drosophila. eLife 6, 1–20. 10.7554/ELIFE.29593.

39. Rossi, M., Hausmann, A.E., Alcami, P., Moest, M., Roussou, R., Van Belleghem, S.M., Wright, D.S., Kuo, C.-Y., Lozano-Urrego, D., Maulana, A., et al. (2024). Adaptive introgression of a visual preference gene. Science 383, 1368–1373. 10.1126/science.adj9201.

40. Kapan, D.D. (2001). Three-butterfly system provides a field test of müllerian mimicry. Nature 409, 338–340. 10.1038/35053066.

41. Chamberlain, N.L., Hill, R.I., Kapan, D.D., Gilbert, L.E., and Kronforst, M.R. (2009). Polymorphic butterfly reveals the missing link in ecological speciation. Science 326, 847–850. 10.1126/science.1179141.

42. Westerman, E.L., VanKuren, N.W., Massardo, D., Tenger-Trolander, A., Zhang, W., Hill, R.I., Perry, M., Bayala, E., Barr, K., Chamberlain, N., et al. (2018). Aristaless controls butterfly wing color variation used in mimicry and mate choice. Curr. Biol. 28, 3469–3474.e4. 10.1016/j.cub.2018.08.051.

43. Manni, M., Berkeley, M.R., Seppey, M., Simão, F.A., and Zdobnov, E.M. (2021). BUSCO update: novel and streamlined workflows along with broader and deeper phylogenetic coverage for scoring of eukaryotic, prokaryotic, and viral genomes. Mol. Biol. Evol. 38, 4647–4654. 10.1093/molbev/msab199.

44. Ruggieri, A.A., Livraghi, L., Lewis, J.J., Evans, E., Cicconardi, F., Hebberecht, L., Ortiz-Ruiz, Y., Montgomery, S.H., Ghezzi, A., Rodriguez-Martinez, J.A., et al. (2022). A butterfly pan-genome reveals that a large amount of structural variation underlies the evolution of chromatin accessibility. Genome Res. 32, 1862–1875. 10.1101/gr.276839.122.

45. Voichek, Y., and Weigel, D. (2020). Identifying genetic variants underlying phenotypic variation in plants without complete genomes. Nat. Genet. 52, 534–540. 10.1038/s41588-020-0612-7.

46. Chen, H., Wang, C., Conomos, M.P., Stilp, A.M., Li, Z., Sofer, T., Szpiro, A.A., Chen, W., Brehm, J.M., Celedón, J.C., et al. (2016). Control for population structure and relatedness for binary traits in genetic association studies via logistic mixed models. Am. J. Hum. Genet. 98, 653–666. 10.1016/j.ajhg.2016.02.012.

47. Merrill, R.M., Rastas, P., Martin, S.H., Melo, M.C., Barker, S., Davey, J., McMillan, W.O., and Jiggins, C.D. (2019). Genetic dissection of assortative mating behavior. PLOS Biol. 17, 1–21. 10.1371/journal.pbio.2005902.

48. Hoffmann, A.A., and Rieseberg, L.H. (2008). Revisiting the impact of inversions in evolution: from population genetic markers to drivers of adaptive shifts and speciation? Annu. Rev. Ecol. Evol. Syst. 39, 21–42. 10.1146/annurev.ecolsys.39.110707.173532.

49. Nowling, R.J., Manke, K.R., and Emrich, S.J. (2020). Detecting inversions with PCA in the presence of population structure. PLOS One 15, e0240429. 10.1371/journal.pone.0240429.

50. Heller, D., and Vingron, M. (2019). SVIM: structural variant identification using mapped long reads. Bioinformatics 35, 2907–2915. 10.1093/bioinformatics/btz041.

51. Davey, J.W., Barker, S.L., Rastas, P.M., Pinharanda, A., Martin, S.H., Durbin, R., McMillan, W.O., Merrill, R.M., and Jiggins, C.D. (2017). No evidence for maintenance of a sympatric heliconius species barrier by chromosomal inversions. Evol. Lett. 1, 138–154. 10.1002/evl3.12.

52. Szabo, Q., Bantignies, F., and Cavalli, G. (2019). Principles of genome folding into topologically associating domains. Sci. Adv. 5, eaaw1668. 10.1126/sciadv.aaw1668.

53. Bonev, B., and Cavalli, G. (2016). Organization and function of the 3D genome. Nat Rev Genet 17, 661–678. 10.1038/nrg.2016.112.

54. Jafar-Nejad, H., and Bellen, H.J. (2004). Gfi/pag-3/senseless zinc finger proteins: a unifying theme? Mol. Cell. Biol. 24, 8803–8812. 10.1128/mcb.24.20.8803-8812.2004.

55. Frankfort, B.J., Pepple, K.L., Mamlouk, M., Rose, M.F., and Mardon, G. (2004). Senseless is required for pupal retinal development inDrosophila. Genesis 38, 182–194. 10.1002/gene.20018.

56. Xie, B., Charlton-Perkins, M., McDonald, E., Gebelein, B., and Cook, T. (2007). Senseless functions as a molecular switch for color photoreceptor differentiation in drosophila. Development 134, 4243–4253. 10.1242/dev.012781.

57. Fisher, W.W., Hammonds, A.S., Weiszmann, R., Booth, B.W., Gevirtzman, L., Patton, J.E.J., Kubo, C.A., Waterston, R.H., Celniker, S.E., Geyer, P., et al. (2023). A modERN resource: identification of *Drosophila* transcription factor candidate target genes using RNAi. Genetics 223, iyad004. 10.1093/genetics/iyad004.

58. Hammonds, A.S., Bristow, C.A., Fisher, W.W., Weiszmann, R., Wu, S., Hartenstein, V., Kellis, M., Yu, B., Frise, E., and Celniker, S.E. (2013). Spatial expression of transcription factors in drosophila embryonic organ development. Genome Biol. 14, R140. 10.1186/gb-2013-14-12-r140.

59. Lacin, H., Zhu, Y., DiPaola, J.T., Wilson, B.A., Zhu, Y., Skeath, J.B., Arbeitman, M., Lacin, H., Zhu, Y., DiPaola, J.T., et al. (2024). A genetic screen in *Drosophila* uncovers a role for *senseless-2* in surface glia in the peripheral nervous system to regulate CNS morphology. G3: Genes, Genomes, Genet. 14, jkae152. 10.1093/g3journal/jkae152.

60. Montell, C., and Rubin, G.M. (1988). The drosophila ninaC locus encodes two photoreceptor cell specific proteins with domains homologous to protein kinases and the myosin heavy chain head. Cell 52, 757–772. 10.1016/0092-8674(88)90413-8.

61. Buenrostro, J.D., Giresi, P.G., Zaba, L.C., Chang, H.Y., and Greenleaf, W.J. (2013). Transposition of native chromatin for fast and sensitive epigenomic profiling of open chromatin, DNA-binding proteins and nucleosome position. Nat Methods 10, 1213–1218. 10.1038/nmeth.2688.

62. Ghavi-Helm, Y., Klein, F.A., Pakozdi, T., Ciglar, L., Noordermeer, D., Huber, W., and Furlong, E.E.M. (2014). Enhancer loops appear stable during development and are associated with paused polymerase. Nature 512, 96–100. 10.1038/nature13417.

63. Lu, W., and Kronforst, M.R. (2025). Cellular innovations and diversity in the lepidopteran compound eye. J. Comp. Physiol. a, 1–20. 10.1007/s00359-025-01751-8.

64. Yildirim, K., Petri, J., Kottmeier, R., and Klämbt, C. (2019). Drosophila glia: few cell types and many conserved functions. Glia 67, 5–26. 10.1002/GLIA.23459.

65. Liu, B., Li, C., Li, Z., Wang, D., Ren, X., Zhang, Z., Liu, B., Li, C., Li, Z., Wang, D., et al. (2020). An entropy-based metric for assessing the purity of single cell populations. Nat. Commun. 11, 3155. 10.1038/s41467-020-16904-3.

66. DiAntonio, A., Burgess, R.W., Chin, A.C., Deitcher, D.L., Scheller, R.H., and Schwarz, T.L. (1993). Identification and characterization of drosophila genes for synaptic vesicle proteins. J. Neurosci. 13, 4924–4935. 10.1523/JNEUROSCI.13-11-04924.1993.

67. Xiong, W.C., Okano, H., Patel, N.H., Blendy, J.A., and Montell, C. (1994). repo encodes a glial-specific homeo domain protein required in the drosophila nervous system. Genes Dev. 8, 981–994. 10.1101/gad.8.8.981.

68. Korsunsky, I., Millard, N., Fan, J., Slowikowski, K., Zhang, F., Wei, K., Baglaenko, Y., Brenner, M., Loh, P., and Raychaudhuri, S. (2019). Fast, sensitive and accurate integration of single-cell data with harmony. Nat. Methods 16, 1289–1296. 10.1038/s41592-019-0619-0.

69. Gao, K., Donati, A., Ainsworth, J., Wu, D., Terner, E.R., and Perry, M.W. (2025). Deep conservation complemented by novelty and innovation in the insect eye ground plan. Proc. Natl. Acad. Sci. 122, e2416562122. 10.1073/pnas.2416562122.

70. Chen, P.-J., Matsushita, A., Wakakuwa, M., and Arikawa, K. (2019). Immunolocalization suggests a role of the histamine-gated chloride channel PxHCLB in spectral opponent processing in butterfly photoreceptors. J. Comp. Neurol. 527, 753–766. 10.1002/cne.24558.

71. Xu, C., Ramos, T.B., Rogers, E.M., Reiser, M.B., and Doe, C.Q. (2023). Homeodomain proteins hierarchically specify neuronal diversity and synaptic connectivity. eLife 12, 1–52. 10.7554/eLife.90133.2.

72. Fu, W., and Noll, M. (1997). The Pax2 homolog sparkling is required for development of cone and pigment cells in theDrosophila eye. Genes Dev. 11, 2066–2078. 10.1101/gad.11.16.2066.

73. Blochlinger, K., Jan, L.Y., and Jan, Y.N. (1993). Postembryonic patterns of expression of cut, a locus regulating sensory organ identity in Drosophila. Development 117, 441–450. 10.1242/dev.117.2.441.

74. Wilk, R., Weizman, I., and Shilo, B.Z. (1996). trachealess encodes a bHLH-PAS protein that is an inducer of tracheal cell fates in drosophila. Genes Dev. 10, 93–102. 10.1101/gad.10.1.93.

75. Tarashansky, A.J., Musser, J.M., Khariton, M., Li, P., Arendt, D., Quake, S.R., and Wang, B. (2021). Mapping single-cell atlases throughout metazoa unravels cell type evolution. eLife 10. 10.7554/ELIFE.66747.

76. Lago-Baldaia, I., Cooper, M., Seroka, A., Trivedi, C., Powell, G.T., Wilson, S.W., Ackerman, S.D., Fernandes, V.M., Smith, C.J., Lago-Baldaia, I., et al. (2023). A drosophila glial cell atlas reveals a mismatch between transcriptional and morphological diversity. PLOS Biol. 21, e3002328. 10.1371/journal.pbio.3002328.

77. Xu, C., Lopez, R., Mehlman, E., Regier, J., Jordan, M.I., and Yosef, N. (2021). Probabilistic harmonization and annotation of single-cell transcriptomics data with deep generative models. Mol. Syst. Biol. 17, MSB20209620. 10.15252/msb.20209620.

78. Rainford, J.L., Hofreiter, M., Nicholson, D.B., and Mayhew, P.J. (2014). Phylogenetic distribution of extant richness suggests metamorphosis is a key innovation driving diversification in insects. PLOS One 9, e109085. 10.1371/journal.pone.0109085.

79. Stavenga, D.G. (2002). Reflections on colourful ommatidia of butterfly eyes. J. Exp. Biol. 205, 1077– 1085. 10.1242/jeb.205.8.1077.

80. Kretzschmar, D., Poeck, B., Roth, H., Ernst, R., Keller, A., Porsch, M., Strauss, R., and Pflugfelder, G.O. (2000). Defective pigment granule biogenesis and aberrant behavior caused by mutations in the drosophila AP-3β adaptin gene ruby. Genetics 155, 213–223. 10.1093/genetics/155.1.213.

81. Stuart, T., Srivastava, A., Madad, S., Lareau, C.A., and Satija, R. (2021). Single-cell chromatin state analysis with signac. Nat. Methods 18, 1333–1341. 10.1038/s41592-021-01282-5.

82. Xie, J., Han, Y., Liang, Y., Peng, L., and Wang, T. Drosophila HisT is a specific histamine transporter that contributes to histamine recycling in glia. Sci. Adv. 8, eabq1780. 10.1126/sciadv.abq1780.

83. Xu, Y., An, F., Borycz, J.A., Borycz, J., Meinertzhagen, I.A., and Wang, T. (2015). Histamine recycling is mediated by CarT, a carcinine transporter in drosophila photoreceptors. PLOS Genet. 11, e1005764. 10.1371/journal.pgen.1005764.

84. Chotard, C., and Salecker, I. (2007). Glial cell development and function in the drosophila visual system. Neuron Glia Biol. 3, 17–25. 10.1017/S1740925X07000592.

85. Kronforst, M.R., Young, L.G., Blume, L.M., and Gilbert, L.E. (2006). Multilocus analyses of admixture and introgression among hybridizing heliconius butterflies. Evol.; Int. J. Org. Evol. 60, 1254–1268. 10.1111/j.0014-3820.2006.tb01203.x.

86. Naisbit, R.E., Jiggins, C.D., Linares, M., Salazar, C., and Mallet, J. (2002). Hybrid sterility, haldane’s rule and speciation in heliconius cydno and H. melpomene. Genetics 161, 1517–1526. 10.1093/genetics/161.4.1517.

87. Mérot, C., Salazar, C., Merrill, R.M., Jiggins, C.D., and Joron, M. (2017). What shapes the continuum of reproductive isolation? Lessons from heliconius butterflies. Proc. R. Soc. B: Biol. Sci. 284, 5–6. 10.1098/rspb.2017.0335.

88. Estrada, C., and Jiggins, C.D. (2002). Patterns of pollen feeding and habitat preference among heliconius species. Ecol. Entomol. 27, 448–456. 10.1046/j.1365-2311.2002.00434.x.

89. Walsh, J.T., Junker, I.P., Chen, Y.-C.D., Chen, Y.-C., Gifford, H., Chen, D.S., and Ding, Y. (2025). High-resolution single-cell analyses reveal evolutionary constraints and evolvability of sexual circuits in drosophila. Proc. Natl. Acad. Sci. 122, 2017. 10.1073/pnas.2516083122.

90. Lee, D., Shahandeh, M.P., Abuin, L., and Benton, R. (2025). Comparative single-cell transcriptomic atlases of drosophilid brains suggest glial evolution during ecological adaptation. PLOS Biol. 23, e3003120. 10.1371/journal.pbio.3003120.

91. Bertram, J., Fulton, B., Tourigny, J.P., Peña-Garcia, Y., Moyle, L.C., and Hahn, M.W. (2023). CAGEE: Computational Analysis of Gene Expression Evolution. Mol Biol Evol 40, msad106. 10.1093/molbev/msad106.

92. Belušič, G., Ilić, M., Meglič, A., and Pirih, P. (2021). Red-green opponency in the long visual fibre photoreceptors of brushfoot butterflies (Nymphalidae). Proceedings of the Royal Society B 288. 10.1098/RSPB.2021.1560.

93. Pirih, P., Ilić, M., Meglič, A., and Belušič, G. (2022). Opponent processing in the retinal mosaic of nymphalid butterflies. Philos. Trans. R. Soc. B: Biol. Sci. 377, 20210275. 10.1098/rstb.2021.0275.

94. Deneris, E.S., and Hobert, O. (2014). Maintenance of postmitotic neuronal cell identity. Nat. Neurosci. 17, 899–907. 10.1038/nn.3731.

95. Mollereau, B., Dominguez, M., Webel, R., Colley, N.J., Keung, B., de Celis, J.F., and Desplan, C. (2001). Two-step process for photoreceptor formation in drosophila. Nature 412, 911–913. 10.1038/35091076.

96. Bollepogu Raja, K.K., Yeung, K., Shim, Y.-K., Li, Y., Chen, R., and Mardon, G. (2023). A single cell genomics atlas of the Drosophila larval eye reveals distinct photoreceptor developmental timelines. Nat Commun 14, 7205. 10.1038/s41467-023-43037-0.

97. Büttner, M., Ostner, J., Müller, C.L., Theis, F.J., and Schubert, B. (2021). scCODA is a bayesian model for compositional single-cell data analysis. Nat. Commun. 12, 6876. 10.1038/s41467-021-27150-6.

98. Kopp, M., Servedio, M.R., Mendelson, T.C., Safran, R.J., Rodríguez, R.L., Hauber, M.E., Scordato, E.C., Symes, L.B., Balakrishnan, C.N., Zonana, D.M., et al. (2018). Mechanisms of assortative mating in speciation with gene flow: connecting theory and empirical research. Am. Nat. 191, 1–20. 10.1086/694889.

99. Lugena, A.B., Goforth, K.M., Shetty, V., Zhang, Y., González-Rodríguez, A., Ramírez, M.I., and Merlin, C. (2026). Seasonal blood-brain barrier plasticity links environmental cues to migratory behavior in monarch butterflies. Preprint at bioRxiv, 10.64898/2026.01.16.699996 https://doi.org/10.64898/2026.01.16.699996.

100. Chaturvedi, R., Reddig, K., and Li, H.-S. (2014). Long-distance mechanism of neurotransmitter recycling mediated by glial network facilitates visual function in drosophila. Proc. Natl. Acad. Sci. 111, 2812–2817. 10.1073/pnas.1323714111.

101. Ilić, M., Chen, P.-J., Pirih, P., Meglič, A., Prevc, J., Yago, M., Belušič, G., and Arikawa, K. (2022). Simple and complex, sexually dimorphic retinal mosaic of fritillary butterflies. Philos. Trans. R. Soc. B: Biol. Sci. 377, 1–8. 10.1098/rstb.2021.0276.

102. Kim, B.Y., Miller, D.E., and Wang, J. (2021). DNA extraction and nanopore library prep from 15-30 whole flies. 1–27. 10.17504/protocols.io.bdfqi3mw.

103. Mohana, G., Dorier, J., Li, X., Mouginot, M., Smith, R.C., Malek, H., Leleu, M., Rodriguez, D., Khadka, J., Rosa, P., et al. (2023). Chromosome-level organization of the regulatory genome in the drosophila nervous system. Cell 186, 3826–3844.e26. 10.1016/j.cell.2023.07.008.

104. Vurture, G.W., Sedlazeck, F.J., Nattestad, M., Underwood, C.J., Fang, H., Gurtowski, J., and Schatz, M.C. (2017). GenomeScope: fast reference-free genome profiling from short reads. Bioinformatics 33, 2202–2204. 10.1093/bioinformatics/btx153.

105. Kolmogorov, M., Yuan, J., Lin, Y., Pevzner, P.A., Kolmogorov, M., Yuan, J., Lin, Y., and Pevzner, P.A. (2019). Assembly of long, error-prone reads using repeat graphs. Nat. Biotechnol. 37, 540–546. 10.1038/s41587-019-0072-8.

106. Guan, D., McCarthy, S.A., Wood, J., Howe, K., Wang, Y., and Durbin, R. (2020). Identifying and removing haplotypic duplication in primary genome assemblies. Bioinformatics 36, 2896–2898. 10.1093/bioinformatics/btaa025.

107. Lorig-Roach, R., Meredith, M., Monlong, J., Jain, M., Olsen, H.E., McNulty, B., Porubsky, D., Montague, T.G., Lucas, J.K., Condon, C., et al. (2024). Phased nanopore assembly with shasta and modular graph phasing with GFAse. Genome Res. 34, 454–468. 10.1101/gr.278268.123.

108. Chakraborty, M., Baldwin-Brown, J.G., Long, A.D., and Emerson, J.J. (2016). Contiguous and accurate de novo assembly of metazoan genomes with modest long read coverage. Nucleic Acids Res. 44, e147. 10.1093/nar/gkw654.

109. Sahara, K., Marec, F., and Traut, W. (1999). TTAGG telomeric repeats in chromosomes of some insects and other arthropods. Chromosome Res.: Int. J. Mol. Supramol. Evol. Asp. Chromosome Biol. 7, 449–460. 10.1023/a:1009297729547.

110. Li, H., and Durbin, R. (2009). Fast and accurate short read alignment with burrows–wheeler transform. Bioinformatics 25, 1754–1760. 10.1093/bioinformatics/btp324.

111. Open2C, Abdennur, N., Fudenberg, G., Flyamer, I.M., Galitsyna, A.A., Goloborodko, A., Imakaev, M., and Venev, S.V. (2024). Pairtools: from sequencing data to chromosome contacts. PLOS Comput. Biol. 20, e1012164. 10.1371/journal.pcbi.1012164.

112. Zhou, C., McCarthy, S.A., Durbin, R., Alkan, C., Zhou, C., McCarthy, S.A., Durbin, R., Alkan, C., Zhou, C., McCarthy, S.A., et al. (2023). YaHS: yet another hi-C scaffolding tool. Bioinformatics 39, btac808. 10.1093/bioinformatics/btac808.

113. Hu, J., Fan, J., Sun, Z., and Liu, S. (2020). NextPolish: a fast and efficient genome polishing tool for long-read assembly. Bioinformatics 36, 2253–2255. 10.1093/bioinformatics/btz891.

114. Li, K., Xu, P., Wang, J., Yi, X., and Jiao, Y. (2023). Identification of errors in draft genome assemblies at single-nucleotide resolution for quality assessment and improvement. Nat. Commun. 14, 6556. 10.1038/s41467-023-42336-w.

115. Dasmahapatra, K.K., Walters, J.R., Briscoe, A.D., Davey, J.W., Whibley, A., Nadeau, N.J., Zimin, A.V., Hughes, D.S.T., Ferguson, L.C., Martin, S.H., et al. (2012). Butterfly genome reveals promiscuous exchange of mimicry adaptations among species. Nature 487, 94–98. 10.1038/nature11041.

116. Jin, J.-J., Yu, W.-B., Yang, J.-B., Song, Y., dePamphilis, C.W., Yi, T.-S., and Li, D.-Z. (2020). GetOrganelle: a fast and versatile toolkit for accurate de novo assembly of organelle genomes. Genome Biol. 21, 241–271. 10.1186/s13059-020-02154-5.

117. Lewis, J.J., and Reed, R.D. (2018). Genome-wide regulatory adaptation shapes population-level genomic landscapes in *Heliconius*. Mol. Biol. Evol., 1–15. 10.1093/molbev/msy209.

118. Lee, J., Christoforo, G., Christoforo, G., Foo, C.S., Probert, C., Kundaje, A., Boley, N., Kohpangwei Dacre, M., and Kim, D. (2016). kundajelab/atac_dnase_pipelines: 0.3.3. (Zenodo). 10.5281/zenodo.211733 https://doi.org/10.5281/zenodo.211733.

119. Li, H. (2018). Minimap2: pairwise alignment for nucleotide sequences. Bioinformatics 34, 3094–3100. 10.1093/bioinformatics/bty191.

120. Kim, D., Langmead, B., and Salzberg, S.L. (2015). HISAT: a fast spliced aligner with low memory requirements. Nat. Methods 12, 357–360. 10.1038/nmeth.3317.

121. Pertea, M., Pertea, G.M., Antonescu, C.M., Chang, T.-C., Mendell, J.T., and Salzberg, S.L. (2015). StringTie enables improved reconstruction of a transcriptome from RNA-seq reads. Nat. Biotechnol. 33, 290–295. 10.1038/nbt.3122.

122. Niknafs, Y.S., Pandian, B., Iyer, H.K., Chinnaiyan, A.M., and Iyer, M.K. (2017). TACO produces robust multisample transcriptome assemblies from RNA-seq. Nat. Methods 14, 68–70. 10.1038/nmeth.4078.

123. Gabriel, L., Brůna, T., Hoff, K.J., Ebel, M., Lomsadze, A., Borodovsky, M., and Stanke, M. (2024). BRAKER3: fully automated genome annotation using RNA-seq and protein evidence with GeneMark-ETP, AUGUSTUS, and TSEBRA. Genome Res. 34, 769–777. 10.1101/gr.278090.123.

124. Tegenfeldt, F., Kuznetsov, D., Manni, M., Berkeley, M., Zdobnov, E.M., and Kriventseva, E.V. (2025). OrthoDB and BUSCO update: annotation of orthologs with wider sampling of genomes. Nucleic Acids Res. 53, D516–D522. 10.1093/nar/gkae987.

125. Buchfink, B., Reuter, K., and Drost, H.-G. (2021). Sensitive protein alignments at tree-of-life scale using DIAMOND. Nat Methods 18, 366–368. 10.1038/s41592-021-01101-x.

126. Gotoh, O. (2024). Spaln3: improvement in speed and accuracy of genome mapping and spliced alignment of protein query sequences. Bioinformatics 40, btae517. 10.1093/bioinformatics/btae517.

127. Brůna, T., Lomsadze, A., and Borodovsky, M. (2024). GeneMark-ETP significantly improves the accuracy of automatic annotation of large eukaryotic genomes. Genome Research 34, 757. 10.1101/gr.278373.123.

128. Stanke, M., Diekhans, M., Baertsch, R., and Haussler, D. (2008). Using native and syntenically mapped cDNA alignments to improve de novo gene finding. Bioinformatics 24, 637–644. 10.1093/bioinformatics/btn013.

129. Gabriel, L., Hoff, K.J., Brůna, T., Borodovsky, M., and Stanke, M. (2021). TSEBRA: transcript selector for BRAKER. BMC Bioinf. 22, 566–577. 10.1186/s12859-021-04482-0.

130. Venturini, L., Caim, S., Kaithakottil, G.G., Mapleson, D.L., and Swarbreck, D. (2018). Leveraging multiple transcriptome assembly methods for improved gene structure annotation. GigaScience 7, giy093. 10.1093/gigascience/giy093.

131. Shumate, A., and Salzberg, S.L. (2021). Liftoff: accurate mapping of gene annotations. Bioinformatics 37, 1639–1643. 10.1093/bioinformatics/btaa1016.

132. Darragh, K., Orteu, A., Black, D., Byers, K.J.R.P., Szczerbowski, D., Warren, I.A., Rastas, P., Pinharanda, A., Davey, J.W., Fernanda Garza, S., et al. (2021). A novel terpene synthase controls differences in anti-aphrodisiac pheromone production between closely related heliconius butterflies. PLOS Biol. 19, e3001022. 10.1371/journal.pbio.3001022.

133. Patro, R., Duggal, G., Love, M.I., Irizarry, R.A., and Kingsford, C. (2017). Salmon provides fast and bias-aware quantification of transcript expression. Nat. Methods 14, 417–419. 10.1038/nmeth.4197.

134. Kang, Y.-J., Yang, D.-C., Kong, L., Hou, M., Meng, Y.-Q., Wei, L., and Gao, G. (2017). CPC2: a fast and accurate coding potential calculator based on sequence intrinsic features. Nucleic Acids Res. 45, W12–W16. 10.1093/nar/gkx428.

135. Donath, A., Jühling, F., Al-Arab, M., Bernhart, S.H., Reinhardt, F., Stadler, P.F., Middendorf, M., and Bernt, M. (2019). Improved annotation of protein-coding genes boundaries in metazoan mitochondrial genomes. Nucleic Acids Res. 47, 10543–10552. 10.1093/nar/gkz833.

136. Emms, D.M., and Kelly, S. (2019). OrthoFinder: phylogenetic orthology inference for comparative genomics. Genome Biol. 20, 238–251. 10.1186/s13059-019-1832-y.

137. Huerta-Cepas, J., Szklarczyk, D., Heller, D., Hernández-Plaza, A., Forslund, S.K., Cook, H., Mende, D.R., Letunic, I., Rattei, T., Jensen, L.J., et al. (2019). eggNOG 5.0: a hierarchical, functionally and phylogenetically annotated orthology resource based on 5090 organisms and 2502 viruses. Nucleic Acids Res. 47, D309–D314. 10.1093/nar/gky1085.

138. Corut, A.K., and Wallace, J.G. (2024). kGWASflow: a modular, flexible, and reproducible snakemake workflow for k-mers-based GWAS. G3 Genes|Genomes|Genet. 14, jkad246. 10.1093/g3journal/jkad246.

139. Kokot, M., Długosz, M., and Deorowicz, S. (2017). KMC 3: counting and manipulating k-mer statistics. Bioinformatics 33, 2759–2761. 10.1093/bioinformatics/btx304.

140. Bushnell, B., and Bushnell, B. (2014). BBMap: a fast, accurate, splice-aware aligner. eScholarsh. (Calif. DIGIT. Libr.), 1–3.

141. McLaughlin, C.N., Qi, Y., Quake, S.R., Luo, L., and Li, H. (2022). Isolation and RNA sequencing of single nuclei from drosophila tissues. STAR Protoc. 3, 101417. 10.1016/j.xpro.2022.101417.

142. Fleming, S.J., Chaffin, M.D., Arduini, A., Akkad, A.-D., Banks, E., Marioni, J.C., Philippakis, A.A., Ellinor, P.T., and Babadi, M. (2023). Unsupervised removal of systematic background noise from droplet-based single-cell experiments using CellBender. Nat. Methods 20, 1323–1335. 10.1038/s41592-023-01943-7.

143. Rogozhnikov, A., Ramkumar, P., Shah, K., Bedi, R., Kato, S., and Escola, G.S. (2021). Demuxalot: scaled up genetic demultiplexing for single-cell sequencing. Preprint at bioRxiv, 10.1101/2021.05.22.443646 https://doi.org/10.1101/2021.05.22.443646.

144. Germain, P.-L., Lun, A., Meixide, C.G., Macnair, W., and Robinson, M.D. (2022). Doublet identification in single-cell sequencing data using *scDblFinder*. Preprint at F1000Research, 10.12688/f1000research.73600.2 https://doi.org/10.12688/f1000research.73600.2.

145. Kuehn, E., Clausen, D.S., Null, R.W., Metzger, B.M., Willis, A.D., and Özpolat, B.D. (2022). Segment number threshold determines juvenile onset of germline cluster expansion in platynereis dumerilii. J. Exp. Zoolog. B Mol. Dev. Evol. 338, 225–240. 10.1002/jez.b.23100.

146. Zhang, C., Nielsen, R., and Mirarab, S. (2025). CASTER: direct species tree inference from whole-genome alignments. Sci. (N. Y. N.Y.) 387, eadk9688. 10.1126/science.adk9688.

147. The Darwin Tree of Life Project Consortium (2022). Sequence locally, think globally: the darwin tree of life project. Proc. Natl. Acad. Sci. U. S. A. 119, e2115642118. 10.1073/pnas.2115642118.

