## Supplemental Figures for "Evolution of compound eye cell types shapes visual behaviors across *Heliconius* butterflies"

### Figures S1–S11

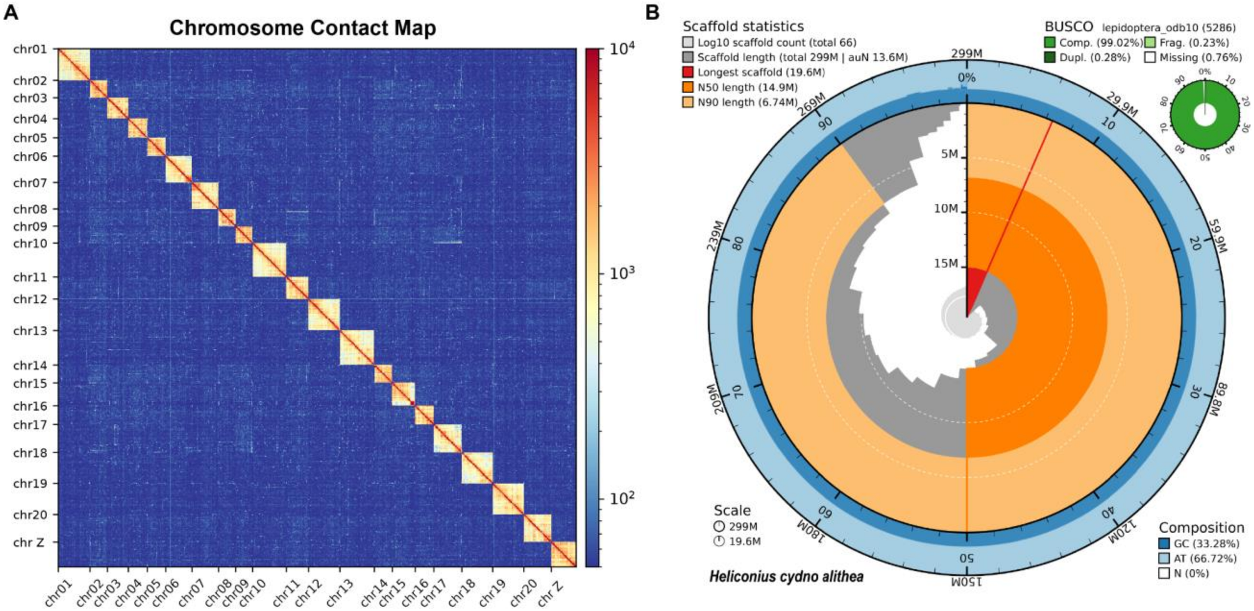


**Figure S1: Genome assembly statistics and contact map for *Heliconius cydno alithea*.** **(A)** Micro-C contact map of the *H. c. alithea* assembly at 200 kb resolution (KR-normalized), illustrating highly contiguous and well-assembled chromosomes. Visualized using HiCExplorer[1]. **(B)** Snail plot summarizing the genome assembly. The circumference represents total genome length, with the outermost blue tracks showing GC/AT content. Scaffolds are ordered clockwise by length (dark grey). Arcs denote the longest scaffold (red), N50 (deep orange), and N90 (pale orange) lengths. The central light grey spiral indicates the cumulative scaffold count on a log10 scale. BUSCO completeness scores are summarized in the top right. Visualized using BlobTools[2].

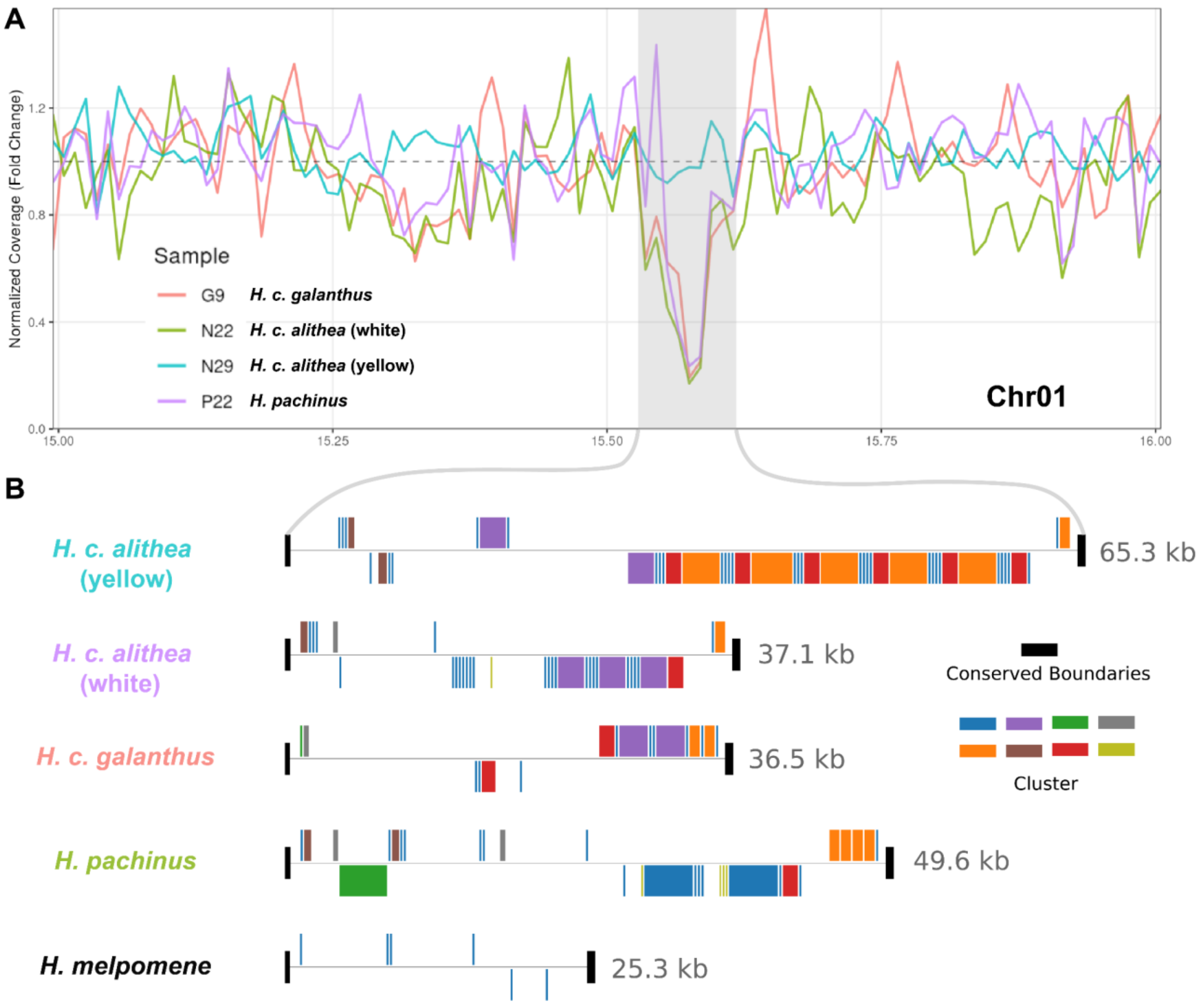


**Figure S2: Structural variation of the 5S rDNA cluster at the *K* locus.**
**(A)** Normalized Nanopore read coverage across the *K* locus on Chromosome 1. Coverage is calculated in 10 kb windows and normalized to average genome-wide read depth (dashed line = 1.0 fold change). Colored lines represent distinct *Heliconius* samples. The gray shaded area highlights a repeat-rich region containing the 5S rDNA cluster, where mapping coverage drops significantly for all individuals except the yellow *H. c. alithea* (N29) sample used to generate the reference assembly. **(B)** *De novo* assemblies resolving the rDNA cluster region across the four individuals. Black rectangles denote conserved DNA boundary sequences anchoring the assemblies. To illustrate structural micro-homology and rearrangement, sequences downstream of the 5S rDNA were grouped by similarity into distinct clusters, represented by the colored blocks. Blocks positioned above the central horizontal line indicate the sequence is on the forward strand, while those below indicate the reverse strand. Total assembled lengths for each region are noted on the right.

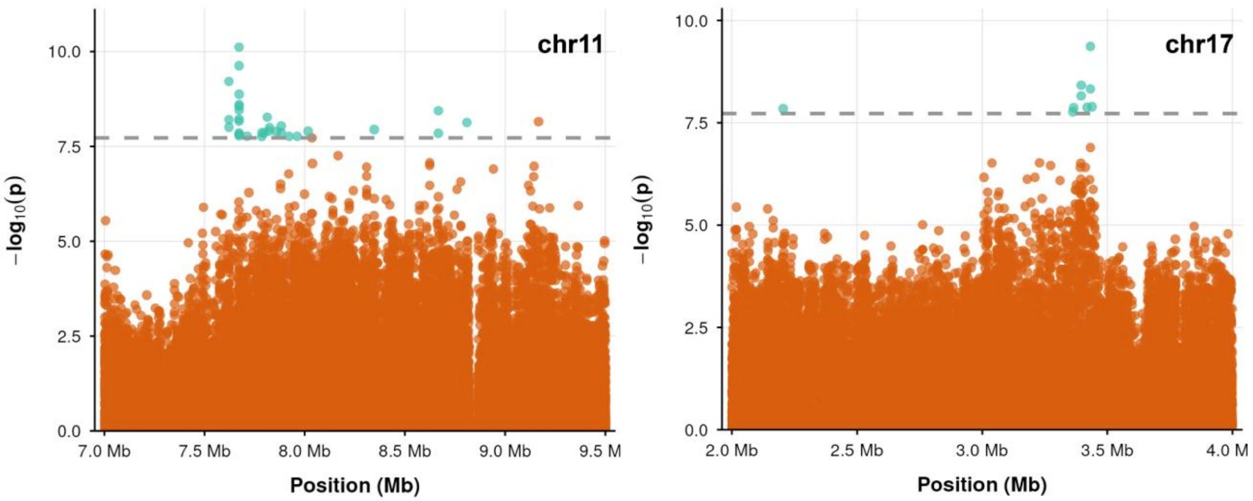


**Figure S3: GWAS peaks outside chromosome 1.**
Zoom-in view of the Manhattan plot across the 2-2.5 Mb region on chromosome 11 and 17. Orange and cyan points represent SNP- and *k*-mer-based associations, respectively. Only *k*-mers passing a false discovery rate (FDR) < 0.01 are displayed. Dash lines indicate the 1% FDR threshold.

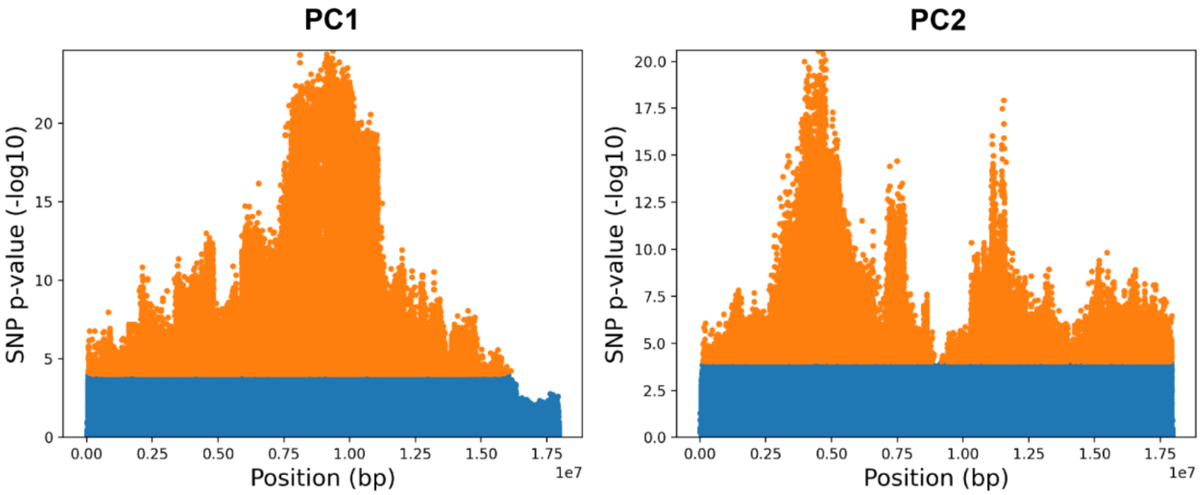


**Figure S4: Absence of a large inversion on chromosome 1.**
Manhattan plots showing SNP associations with PC1 and PC2 for 109 *H. c. alithea* individuals, generated using Asaph[3]. The absence of a step-function pattern indicates a lack of a large structural inversion, particularly within the 15–16 Mb region containing the *K* locus. SNPs highlighted in orange denote significant associations at a 0.01 threshold following Bonferroni correction.

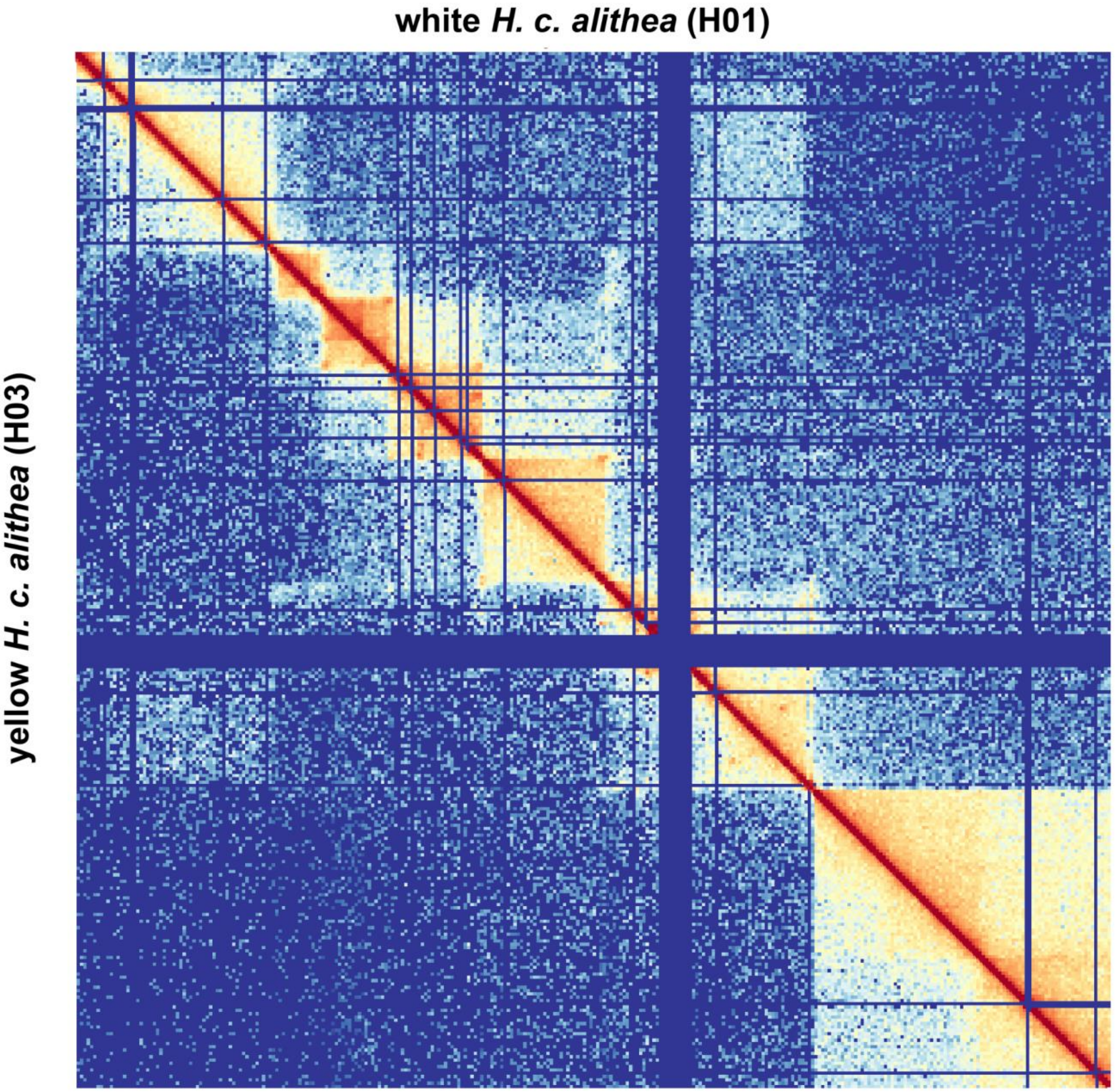


**Figure S5: Conserved 3D chromatin architecture at the K locus between *H. c. alithea* morphs.**
Composite ICE-normalized Micro-C contact map of chromosome 1 (15–16 Mb). Bin size is 3.2 kb. Data from the white male library (H01) is plotted on the upper triangle, while data from the yellow male library (H03) is plotted on the lower triangle. The loops, TADs, and compartments of the *K* locus remain consistent between the two color morphs.

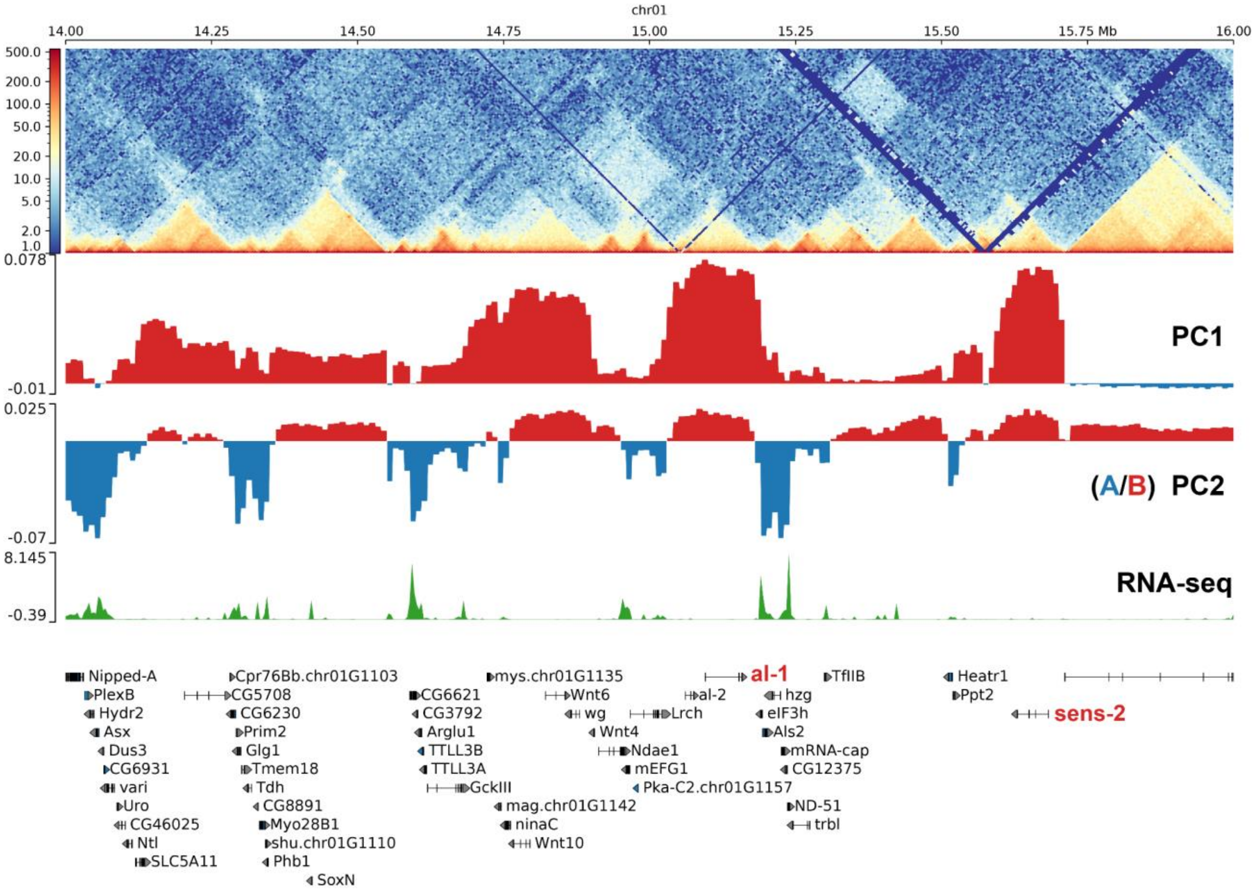


**Figure S6: A/B compartmentalization at the K locus.**

Top to bottom: Micro-C contact map at 5 kb resolution spanning 14–16 Mb on chromosome 1; Principal Component 1 (PC1) and 2 (PC2) eigenvalues derived from the 10 kb KR-normalized Micro-C matrix. In this chromosome, PC2 delineates the chromatin compartments, where negative values indicate active (A) compartments and positive values indicate inactive (B) compartments. Active compartments correlate with regions of higher gene density and elevated gene expression, as demonstrated by the adult optic lobe RNA-seq track. The key color and color-preference genes, *al-1* and *sens-2* (highlighted in red), both localize to the inactive B compartment in adult brain tissue.

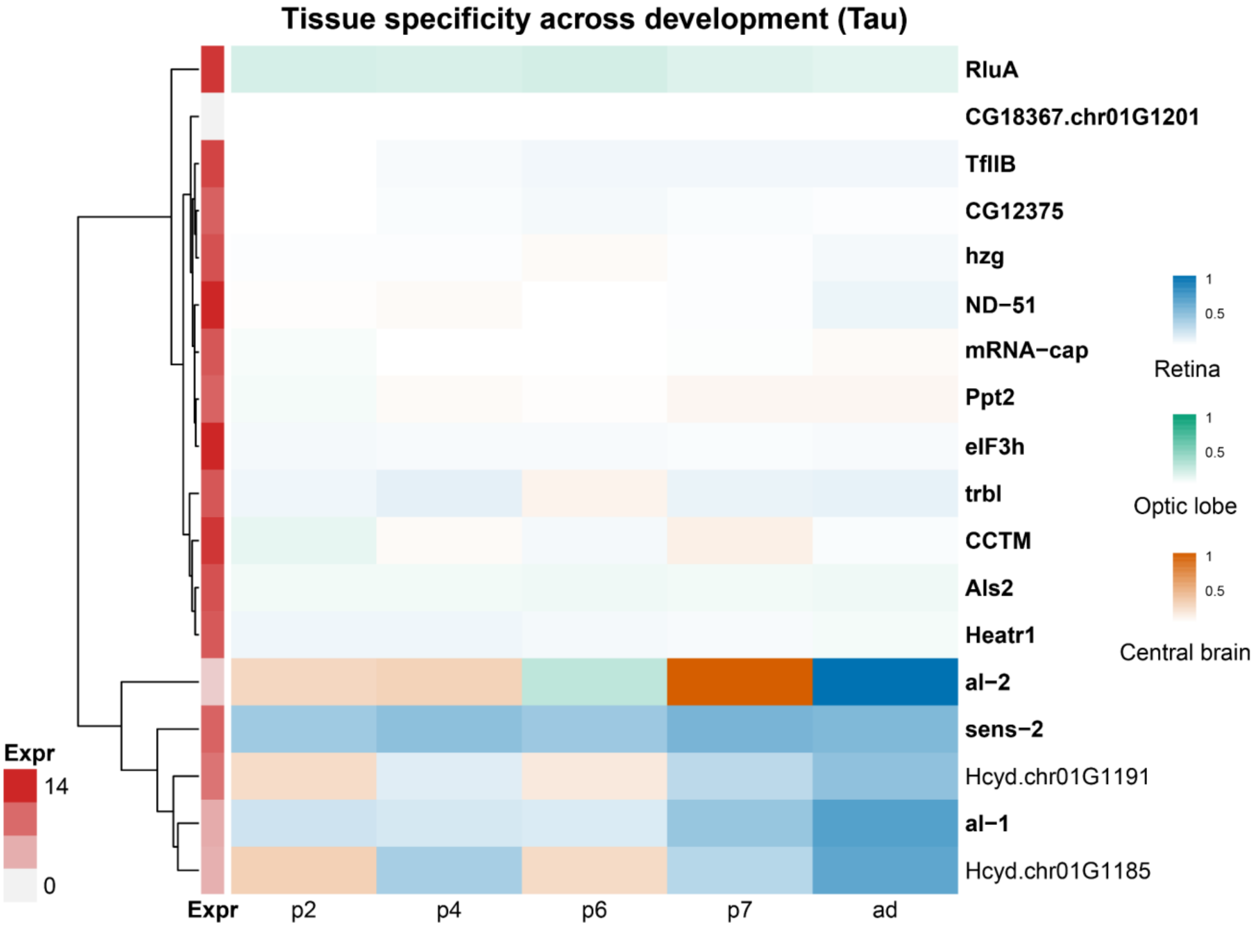


**Figure S7: Tissue specificity of K locus genes across visual system development.** Heatmap displaying the tissue specificity of candidate genes across pupal stages (p2, p4, p6, p7) and adult (ad). Specificity was calculated using the Tau index on log2-normalized bulk RNA-seq counts. The color hue indicates the tissue with peak expression, while color saturation represents the magnitude of the Tau index, with darker shades indicating stronger tissue specificity (closer to 1). The leftmost annotation (Expr) denotes the maximum expression level across stages and tissues for each gene. Rows are ordered via hierarchical clustering. Many housekeeping genes, such as *eIF3h* and *TfIIB*, cluster together and show low tissue specificity. Notably, the candidate color-preference gene *sens-2* demonstrates a strictly retina-specific expression pattern throughout all stages of pupal development and into the adult.

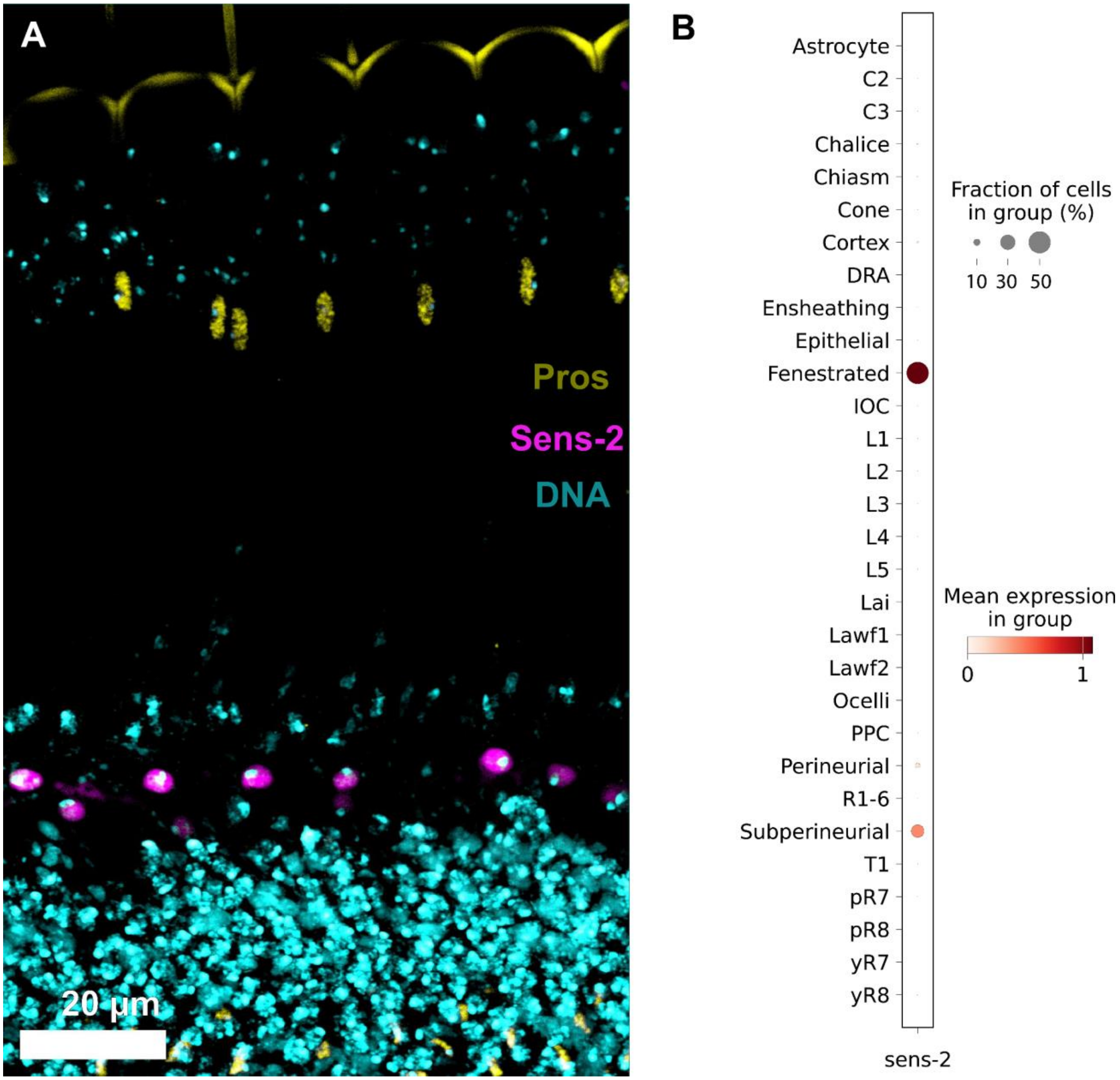


**Figure S8: *sens-2* expression pattern in the *Drosophila* visual system.**
**(A)** Confocal cross-section of the *Drosophila* adult compound eye. Tissue is stained for Prospero (Pros, yellow) to indicate R7 photoreceptor positions, Sens-2 (magenta), and DNA (cyan). *sens-2* expression is restricted to the basal fenestrated glia layer at the boundary between the retina and the optic lobe. Scale bar is 20 µm. **(B)** Dot plot displaying *sens-2* transcript expression across visual system cell types derived from the *Drosophila* Fly Cell Atlas (FCA) eye snRNA-seq dataset. Consistent with the staining result, *sens-2* expression is highly enriched in fenestrated and subperineurial glia.

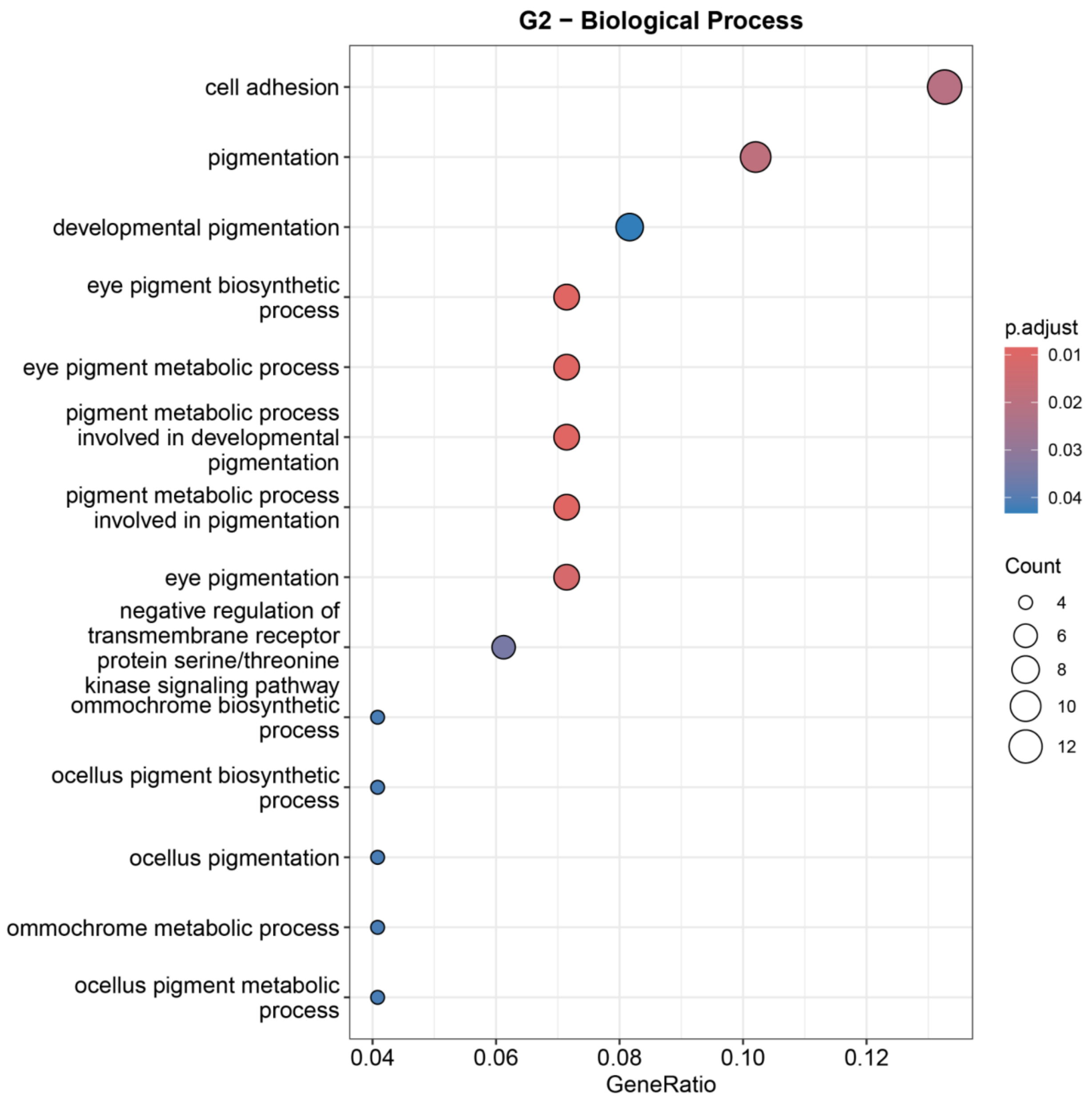


**Figure S9: Gene Ontology (GO) Biological Process enrichment for the G2 cell cluster.** Dot plot displaying the significantly enriched GO biological process terms for the top marker genes of the G2 cell cluster. The x-axis represents the proportion of input marker genes associated with a specific GO term. Node color indicates the Benjamini-Hochberg-adjusted p-value. Node size corresponds to the number of genes mapped to each term. The G2 cluster exhibits strong and highly significant enrichment for pigmentation-related GO terms, specifically eye pigment and ommochrome biosynthetic processes, indicating a pigment cell identity.

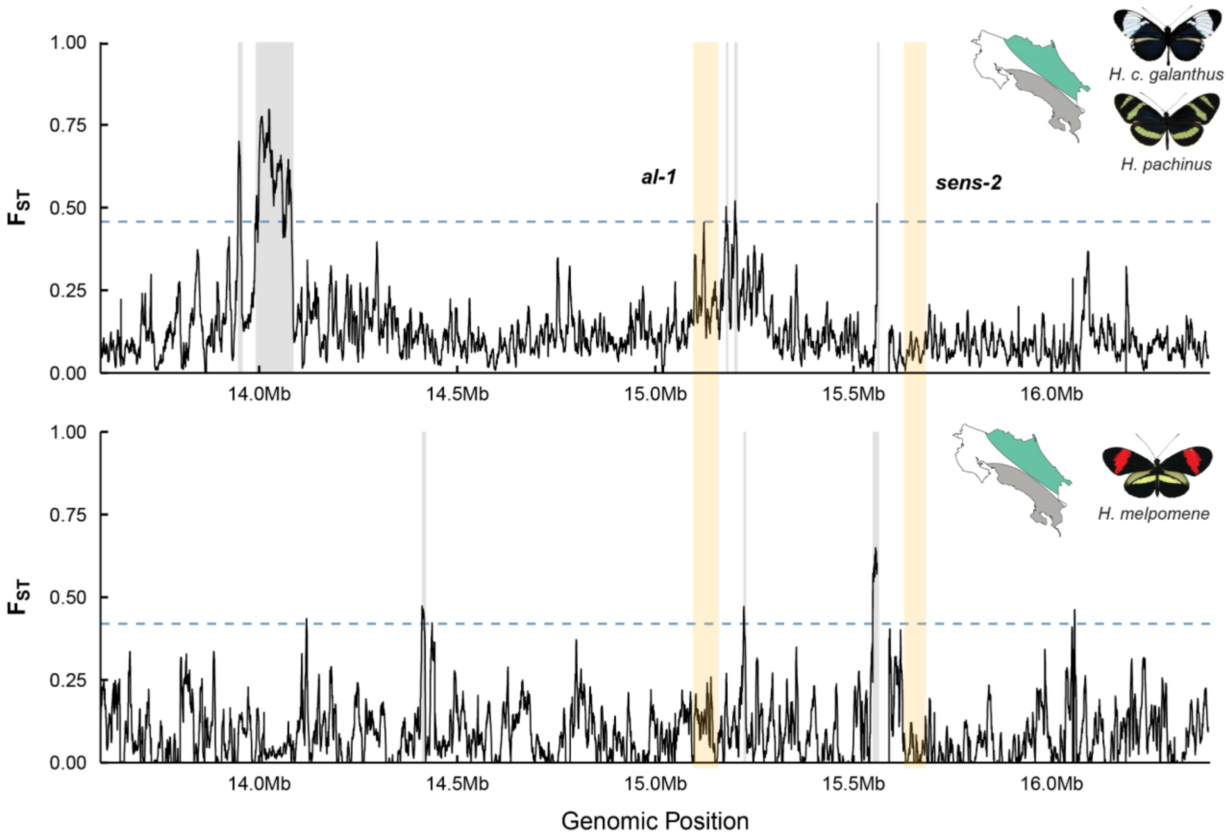


**Figure S10: Sliding window $F_{st}$ across the broader K locus region.**
**(Top panel)** $F_{st}$ between parapatric *H. c. galanthus* and *H. pachinus* in Costa Rica (ranges indicated on the inset map). **(Bottom panel)** $F_{st}$ between Caribbean and Pacific populations of *H. melpomene*. Yellow shaded bars denote genes of interest, *al-1* and *sens-2*. The dashed blue line indicates the threshold for the top 0.5% of $F_{st}$ windows. $F_{st}$ windows above the 0.5% threshold are colored in gray-shaded bars. $F_{st}$ was calculated in 5-kb windows with a 200-bp step size.

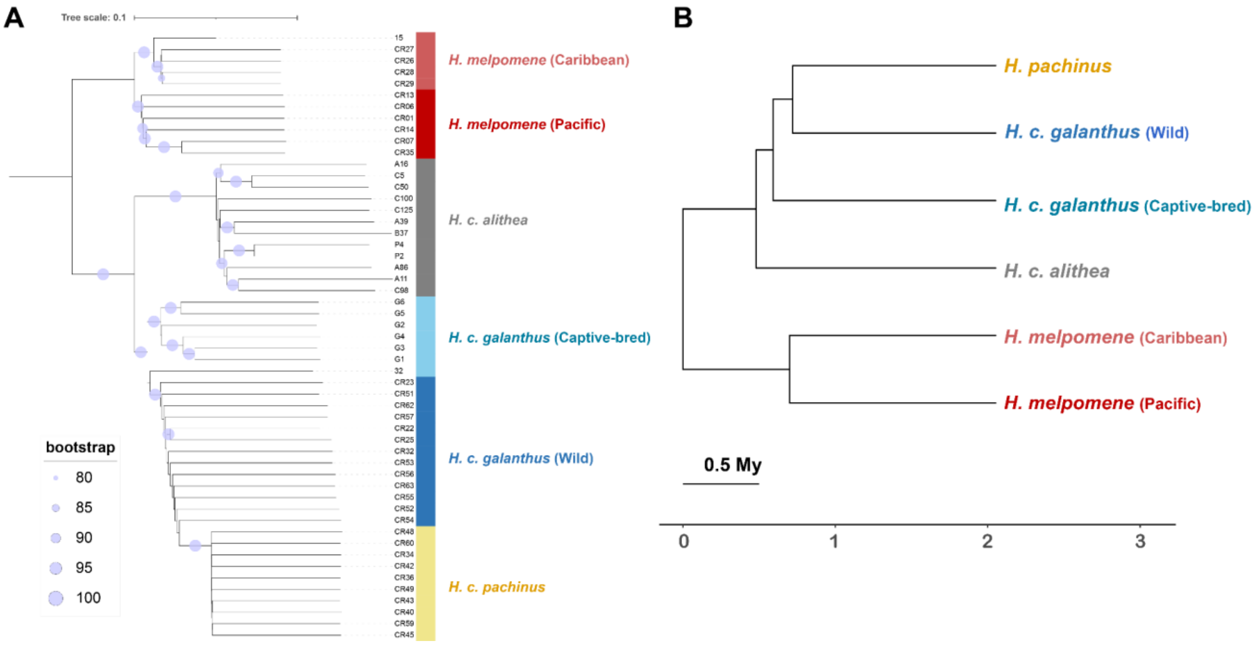


**Figure S11: Phylogeny and divergence times of sequenced *Heliconius* populations. (A)** Phylogenetic tree of sequenced *Heliconius* individuals. Node support is indicated by light purple circles, with circle size proportional to bootstrap values. The scale bar (top left) represents 0.1 substitutions per site. The tree resolves the relationships between *H. melpomene* (Caribbean and Pacific populations) and the *H. cydno* complex, including *H. c. alithea*, paraphyletic *H. c. galanthus*, and *H. pachinus*. **(B)** Time-calibrated population tree estimating divergence times among the represented population. The x-axis and scale bar indicate evolutionary time in millions of years (My).

### References


1. Ramírez, F., Bhardwaj, V., Arrigoni, L., Lam, K.C., Grüning, B.A., Villaveces, J., Habermann, B., Akhtar, A., and Manke, T. (2018). High-resolution TADs reveal DNA sequences underlying genome organization in flies. Nat. Commun. *9*, 189–203. https://doi.org/10.1038/s41467-017-02525-w.

2. Challis, R., Richards, E., Rajan, J., Cochrane, G., and Blaxter, M. (2020). BlobToolKit – interactive quality assessment of genome assemblies. G3: Genes|Genomes|Genet. *10*, 1361–1374. https://doi.org/10.1534/g3.119.400908.

3. Nowling, R.J., Manke, K.R., and Emrich, S.J. (2020). Detecting inversions with PCA in the presence of population structure. PLOS One *15*, e0240429. https://doi.org/10.1371/journal.pone.0240429.
